# Chemo-senolytic therapeutic potential against angiosarcoma

**DOI:** 10.1101/2023.10.19.563192

**Authors:** Xuebing Wang, Claire Yik-Lok Chung, Ai Yoshioka, Shinya Hashimoto, Haruki Jimbo, Hideki Tanizawa, Shinya Ohta, Takeshi Fukumoto, Ken-ichi Noma

## Abstract

Angiosarcoma is an aggressive soft-tissue sarcoma with a poor prognosis. Chemotherapy for this cancer typically employs paclitaxel, one of the taxanes (genotoxic drugs), although it has a limited effect due to chemoresistance for prolonged treatment. Here we examine a new angiosarcoma treatment approach that combines chemotherapeutic and senolytic agents. We first find that the chemotherapeutic drugs, cisplatin and paclitaxel, efficiently induce cellular senescence of angiosarcoma cells. Subsequent treatment with a senolytic agent, ABT-263, eliminates senescent cells through the activation of the apoptotic pathway. In addition, expression analysis indicates that senescence-associated secretory phenotype (SASP) genes are activated in senescent angiosarcoma cells and that ABT-263 treatment eliminates senescent cells expressing genes in the type-I interferon (IFN-I) pathway. Moreover, we show that cisplatin treatment alone requires a high dose to remove angiosarcoma cells, whereas a lower dose of cisplatin is sufficient to induce senescence, followed by the elimination of senescent cells by senolytic treatment. This study sheds light on a potential therapeutic strategy against angiosarcoma by combining a relatively low dose of cisplatin with the ABT-263 senolytic agent, which can help ease the deleterious side effects of chemotherapy.

## Introduction

Angiosarcoma is an aggressive soft-tissue sarcoma that arises from vascular or lymphoid endothelial cells, with an incidence of approximately 1-3 cases per 1,000,000 people per year and a 5-year survival rate of approximately 10-50% (Abraham et al., 2007, Albores-Saavedra et al., 2011, Bi et al., 2022, Cioffi et al., 2013, Lee et al., 2019, Young et al., 2010). Treatment of angiosarcoma remains challenging due to limited evidence with randomized controlled phase 3 clinical trials and few prospective phase 2 studies (Young et al., 2010). Currently, for patients in resectable stages, surgery is the first choice. However, it has been reported that the risk of recurrence remains high in patients who undergo complete resection (Lahat et al., 2010). Therefore, chemotherapy is used to prevent postoperative recurrence and when surgical resection is difficult (Young et al., 2010).

Taxanes are frequently used in single-agent chemotherapy for treatment of angiosarcoma (Penel et al., 2008, Young et al., 2010). Paclitaxel and docetaxel, the taxane genotoxic drugs recommended by the National Comprehensive Cancer Network (NCCN), have been reported to be effective for various types of angiosarcoma, including advanced head angiosarcoma (Fata et al., 1999, Schlemmer et al., 2008). Paclitaxel is the most commonly used chemotherapeutic drug for treatment of angiosarcoma, but its long-term inhibitory effect on tumors remains limited. For instance, a study shows that the median time to progression of paclitaxel treatment is 7.6 months for patients with angiosarcoma, which can partly explain the high risk of recurrence (Schlemmer et al., 2008). Despite the accumulation of insights into angiosarcoma treatment, a key hindrance still exists for current strategies, and it stems from the recurrence of angiosarcoma due to drug resistance, although systemic treatment initially works as expected. Therefore, developing an alternative strategy is a critical challenge to prolong the survival of patients with angiosarcoma.

Beyond angiosarcoma, genotoxic chemotherapeutic drugs, including cisplatin, bleomycin, and doxorubicin, are used to treat various cancers (Chen and Stubbe, 2005, Gabizon et al., 2016, Quinn et al., 2017, Rottenberg et al., 2021). Treatment with genotoxic drugs initiates a DNA damage response, which leads to apoptosis (programmed cell death) if DNA damages are irreparable or cellular senescence (stable cell-cycle arrest) (Hernandez-Segura et al., 2018, Munoz-Espin and Serrano, 2014). Senescent cells are known to produce senescence-associated secretory phenotype (SASP) factors, which are composed of a variety of pro-inflammatory factors and have pleiotropic effects on cancer (Gorgoulis et al., 2019). In addition, recent studies have shown that senescent cells derived from treatment with chemotherapeutic drugs can escape cell-cycle arrest and re-proliferate, at least partially explaining cancer relapse after chemotherapy (Duy et al., 2021, Milanovic et al., 2018, Saleh et al., 2019). Therefore, a one-two punch (chemo-senolytic) approach, in which cancer cells become senescent by chemotherapeutic drug treatment and senescent cancer cells are subsequently eliminated by senolytic treatment, is advantageous for cancer therapy (Di Micco et al., 2021, Paez-Ribes et al., 2019, Wang et al., 2022).

Speaking of senolytics, pharmacological removal of senescent cells used to be difficult until senolytics have recently been identified (Paez-Ribes et al., 2019, Zhu et al., 2015). For instance, the Bcl-2 inhibitor, ABT-263, can remove senescent cells in humans and mice (Chang et al., 2016, Zhu et al., 2016). Importantly, ABT-263 can eliminate senescent cells derived from cancer cells, referred to as senescent cancer cells (Demaria et al., 2017, Jochems et al., 2021, Wang et al., 2017). Although ABT-263 is reported to have platelet toxicity, recent advances have been made to reduce the toxicity of ABT-263 by conjugating it with cleavable galactose or by encapsulating it with galacto-oligosaccharides, allowing this senolytic agent to be preferentially activated in senescent cells (Gonzalez-Gualda et al., 2020, Munoz-Espin et al., 2018). Moreover, there are other types of senolytic agents, such as BPTES (inhibitor of glutaminase 1) and RSL3 (inhibitor of GPX4 ferroptosis regulator) (Johmura et al., 2021, Liao et al., 2022). However, the optimal combination of chemo-senolytic agents remains to be explored for respective cancers. Importantly, the efficacy of the chemo-senolytic approach against angiosarcoma is literally unknown.

Therefore, here we determine the efficacy of the chemo-senolytic treatment for angiosarcoma and the effective combination of chemotherapeutic and senolytic agents for angiosarcoma treatment. We first describe that the two chemotherapy drugs, paclitaxel and cisplatin, can induce cellular senescence of the MO-LAS-B and ISO-HAS-B human angiosarcoma cell lines. Subsequently, senescent cells are treated with one of the three senolytic agents, ABT-263, BPTES, and RSL3, to remove senescent cells. Among the combinations we tested, the pair of cisplatin and ABT-263 is the most effective against angiosarcoma cells, although paclitaxel is the first-line choice for chemotherapeutic treatment of angiosarcoma. Moreover, our expression study shows that chemotherapeutic drug treatment converts angiosarcoma cells to senescent cells expressing SASP genes, and subsequent ABT-263 senolytic treatment removes senescent angiosarcoma cells expressing key senescence genes in the type-I interferon (IFN-I) pathway.

## Results

### Sensitivity of angiosarcoma cells to cisplatin and paclitaxel

Although chemotherapeutic drug treatment can often induce senescence of cancer cells in response to genotoxic stress, it remains unclear whether and how angiosarcoma cells react to drug treatment and which drugs effectively elicit senescence (Ewald et al., 2010). Therefore, we first investigated the senescence induction of angiosarcoma cells using two chemotherapeutic drugs, cisplatin and paclitaxel; cisplatin is a typical genotoxic drug employed for chemotherapy, and paclitaxel is the most used chemotherapeutic drug for angiosarcoma treatment (Jordan and Wilson, 2004, Painter et al., 2020, Rottenberg et al., 2021). Additionally, to examine the cellular reaction to these drugs, we employed the two angiosarcoma cell lines: MO-LAS-B lymphangiosarcoma cell line (Masuzawa et al., 2012); ISO-HAS-B hemangiosarcoma cell line (Masuzawa et al., 1999).

We observed that increasing cisplatin concentration impaired colony formation of both MO-LAS-B and ISO-HAS-B cells, indicating that cisplatin treatment inhibits the proliferation of angiosarcoma cells (**Figures 1a** and **S1a**). Moreover, we performed cell viability assays and estimated the half maximal inhibitory concentration (IC50) of cisplatin as 7.66 and 38.0 μM for MO-LAS-B and ISO-HAS-B cells, respectively (**Figure 1b-c**). MO-LAS-B lymphangiosarcoma cells were significantly more sensitive to cisplatin than ISO-HAS-B hemangiosarcoma cells. In this regard, ISO-HAS-B cells were known to have mutated *TP53* and elevated expression of some oncogenes compared to MO-LAS-B cells, suggesting that ISO-HAS-B cells are likely derived from a more advanced stage of carcinogenesis compared to MO-LAS-B cells and thus relatively resistant to drug treatment (Masuzawa et al., 1999, Masuzawa et al., 2012).

**Figure 1.**
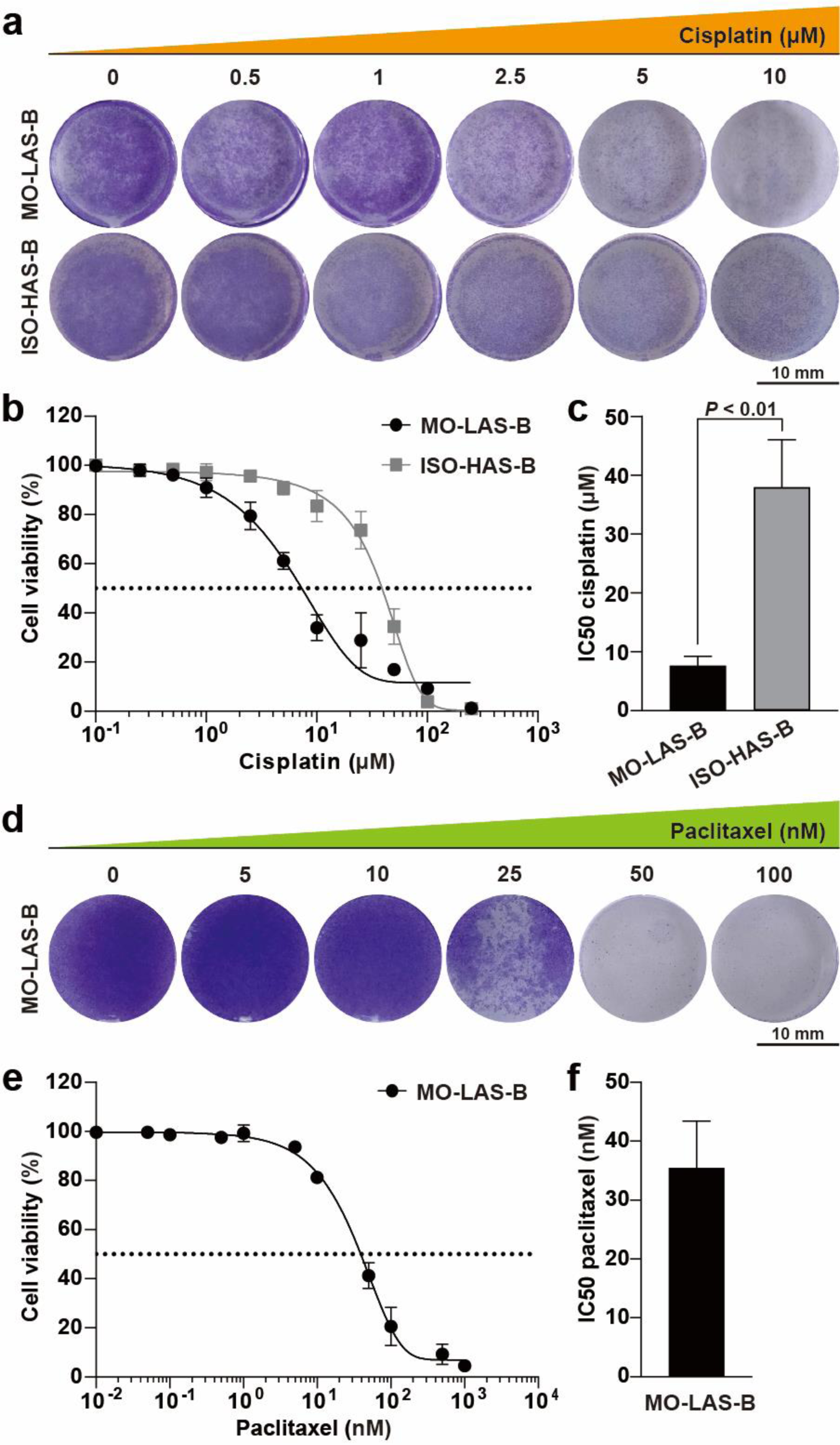
Cisplatin and paclitaxel inhibit proliferation of angiosarcoma cells. **(a)** Effect of cisplatin treatment on colony formation of the angiosarcoma cell lines. MO-LAS-B and ISO-HAS-B cells were treated with cisplatin at the indicated concentrations for 6 days and examined for colony formation. Quantification of the data is shown in **Figure S1a**. **(b)** Cisplatin dose-response curves for MO-LAS-B and ISO-HAS-B cells. Cell numbers after 6 days of culture at the respective cisplatin concentrations were measured, and average values and standard deviations from three biological replicates were plotted. **(c)** Cisplatin IC50 values derived from the dose-response curves in **panel b**. *P*-values were calculated by two-sided Student’s *t*-test, using biologically independent samples (n =3). Error bars represent the standard deviation. **(d)** Effect of paclitaxel treatment on colony formation of the angiosarcoma cell line. MO-LAS-B cells were treated with paclitaxel at the indicated concentrations for 6 days and examined for colony formation. Quantification of the data is shown in **Figure S1b**. **(e)** Paclitaxel dose-response curve for MO-LAS-B cells. Cell numbers after 6 days of culture at the respective paclitaxel concentrations were measured, and the result was plotted as described in **panel b**. **(f)** Paclitaxel IC50 value derived from the dose-response curve in **panel e**.

Moreover, we observed that increasing paclitaxel concentration inhibited colony formation of MO-LAS-B cells (**Figures 1d** and **S1b**); the paclitaxel IC50 value for MO-LAS-B cells was 35.5 nM (**Figure 1e-f**). These results show that the angiosarcoma cell lines we examined have some degree of vulnerability to the chemotherapeutic drugs, although the degree varies depending on cell types and drugs. According to the Genomics of Drug Sensitivity in Cancer (GDSC) database (Yang et al., 2013), the geometric mean of cisplatin IC50 values from 760 cancer cell lines was 25.4 μM, and that of paclitaxel IC50 values from 958 cancer cell lines was 57.5 nM. The IC50 values of cisplatin and paclitaxel were in a similar range for angiosarcoma cells, but the IC50 values were lower for MO-LAS-B cells, suggesting that this cell line is derived from a relatively early stage of carcinogenesis compared to the ISO-HAS-B and many cancer cell lines.

### Senescence induction of angiosarcoma cells by cisplatin

We investigated whether the growth inhibition observed with chemotherapeutic drug treatment was due to the induction of senescence or apoptosis pathways. To that end, we treated angiosarcoma cells with various concentrations of cisplatin and monitored the senescence marker, senescence-associated β-galactosidase (SA-β-gal) activity. SA-β-gal-positive cells were readily detected when cells were treated with cisplatin at the 2.5 and 5 μM concentrations, whereas SA-β-gal-positive cells were almost absent at cisplatin concentrations below 2.5 μM and above 5 μM, suggesting that senescence of angiosarcoma cells can be efficiently induced by cisplatin treatment within a certain concentration range (**Figure 2a**).

**Figure 2.**
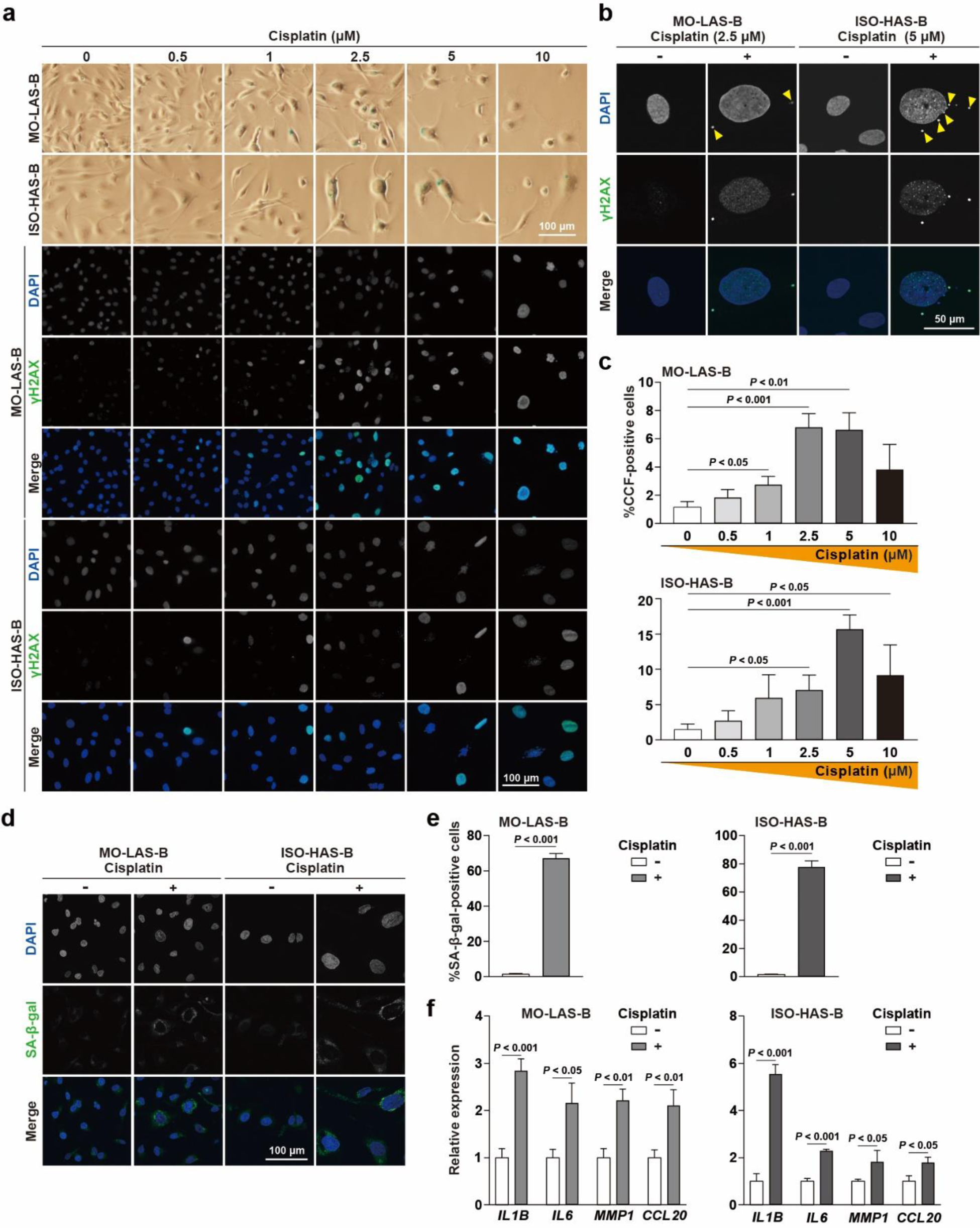
Cisplatin induces senescence of angiosarcoma cells. **(a)** Effect of cisplatin treatment on senescence induction. Cells cultured for 6 days in media containing cisplatin at the indicated concentrations were subjected to SA-β-gal staining (top two rows) and IF experiments to visualize DAPI and γH2AX signals (bottom rows). Quantification of the data is shown in **Figure S2a**. **(b)** CCF detection after cisplatin treatment. DAPI and γH2AX signals were visualized in MO-LAS-B (left) and ISO-HAS-B cells (right) treated with 2.5 and 5 μM cisplatin, respectively (cisplatin +). Untread cells (cisplatin -) are shown as a control. Arrowheads in the DAPI images represent potential CCF. **(c)** Percentages of CCF-positive cells after cisplatin treatment. As shown in **panel b**, cells cultured at the indicated cisplatin concentrations were subjected to IF experiments to visualize DAPI and γH2AX. Cells showing co-localization of cytoplasmic DAPI and γH2AX signals near nuclei were counted as CCF-positive cells. **(d)** SA-β-gal signals in MO-LAS-B (left) and ISO-HAS-B cells (right) treated with 2.5 and 5 μM cisplatin, respectively (cisplatin +). Untread cells (cisplatin -) were also subjected to the same experiment. SA-β-gal signals were visualized by SPiDER-βGal. Flow cytometric data analysis is described in **Figure S2b**. **(e)** Percentages of SA-β-gal-positive cells treated or untread with cisplatin. SA-β-gal-positive cells were estimated, as explained in **Figure S2b**. **(f)** Up-regulation of SASP genes in MO-LAS-B (left) and ISO-HAS-B cells (right) treated with 2.5 and 5 μM cisplatin, respectively. Cells cultured in the presence and absence of cisplatin were examined by RT-qPCR.

It has been reported that DNA damage caused by genotoxic drugs induces senescence, resulting in γH2AX staining in the nucleus and cytoplasmic chromatin fragments (CCF) (Dou et al., 2017, Ivanov et al., 2013). We, therefore, visualized γH2AX foci in cisplatin-treated cells and observed that more γH2AX-positive cells appeared as cisplatin concentration increased (**Figures 2a** and **S2a**). In addition, we observed a significant increase in CCF-positive cells as cisplatin concentration elevated (**Figure 2b-c**). On the other hand, CCF-positive cells declined when cells were treated with 10 μM cisplatin, which was the highest concentration we tested for both cell lines (**Figure 2c**). These results suggest that 2.5 and 5 μM cisplatin are the lowest concentrations that most frequently induce senescence for MO-LAS-B and ISO-HAS-B cells, respectively. Interestingly, SA-β-gal- and CCF-positive cells peaked at the 2.5 or 5 μM cisplatin concentrations (**Figure 2a** and **2c**), whereas γH2AX-positive cells were most abundant at the 10 μM cisplatin concentration (**Figure S2a**), implying that 10 μM cisplatin tends to induce apoptosis rather than senescence.

We next took a relatively objective approach, in which cells were stained by SPiDER-βGal and subjected to flow cytometric analysis (Doura et al., 2016). SA-β-gal signals were detected in angiosarcoma cells treated with cisplatin, and quantification of flow cytometry data showed that cisplatin treatment significantly increased SA-β-gal-positive cells from 1.4 to 67% and 1.4 to 78% for MO-LAS-B and ISO-HAS-B cells, respectively (**Figures 2d-e** and **S2b**). In addition, SASP genes were up-regulated by cisplatin treatment (**Figure 2f**). These results suggest that 2.5 and 5 μM cisplatin are sufficient to efficiently induce senescence of MO-LAS-B and ISO-HAS-B angiosarcoma cells, respectively. Hereinafter, senescent cells derived from angiosarcoma cells are referred to as senescent angiosarcoma cells.

### Paclitaxel-induced senescence of angiosarcoma cells

In addition to cisplatin, we investigated whether paclitaxel could also trigger senescence of angiosarcoma cells and observed that paclitaxel treatment indeed increased SA-β-gal-positive cells (**Figure 3a**). γH2AX-positive nuclei and CCF-positive cells were also increased after paclitaxel treatment (**Figure 3a-c**). Moreover, SA-β-gal-positive cells were quantified by flow cytometry analysis, and we observed that SA-β-gal-positive cells significantly increased roughly from 5.2 to 48% by paclitaxel treatment (**Figures 3d-e** and **S3**). In addition, SASP genes were up-regulated by paclitaxel treatment (**Figure 3f**). These data collectively indicate that paclitaxel can also induce senescence of angiosarcoma cells.

**Figure 3.**
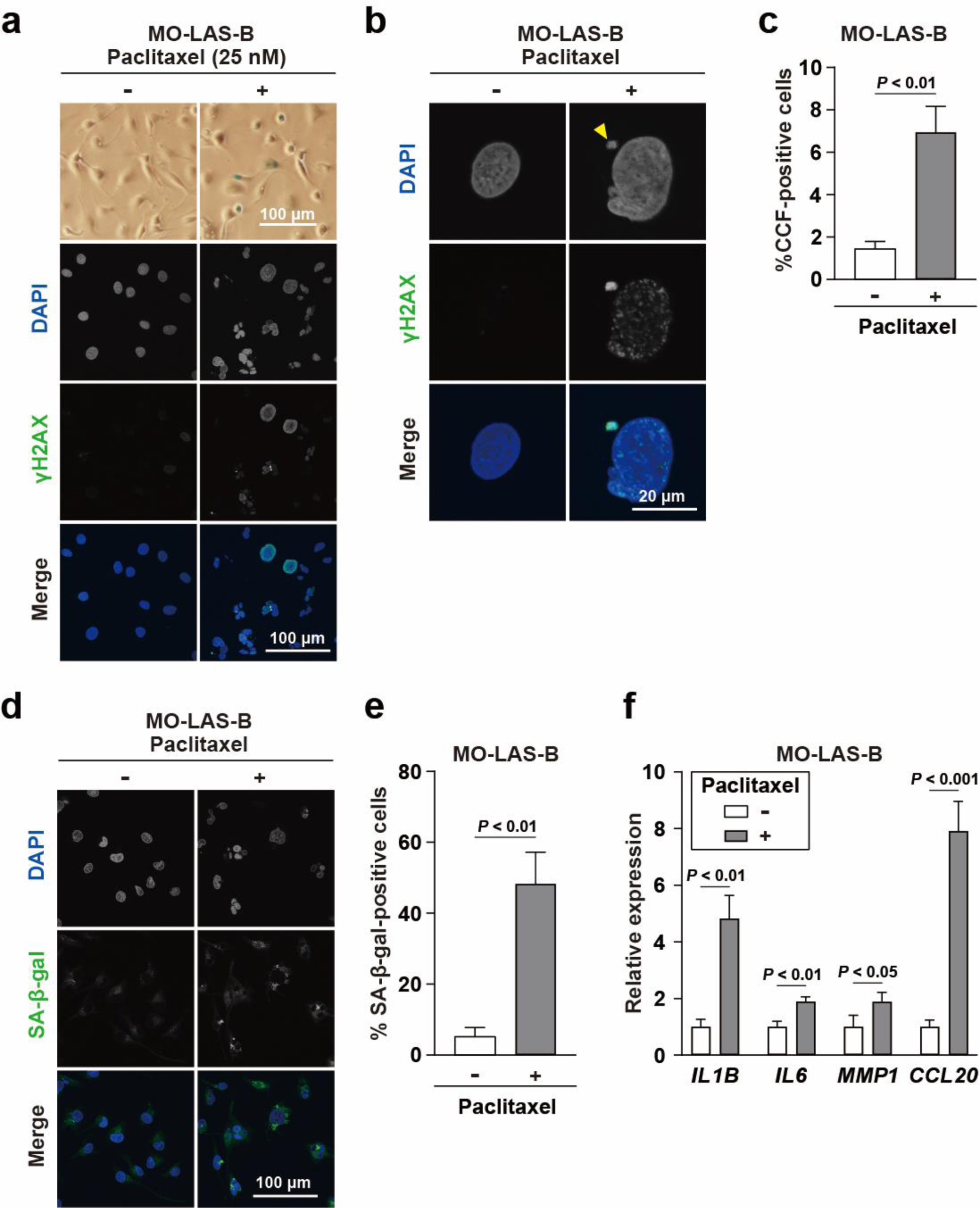
Paclitaxel induces senescence of angiosarcoma cells. **(a)** Effect of paclitaxel treatment on senescence induction. Cells cultured for 6 days in a medium containing 25 nM paclitaxel were subjected to SA-β-gal staining (top row) and IF experiments to visualize DAPI and γH2AX signals (bottom three rows). **(b)** CCF detection after paclitaxel treatment. DAPI and γH2AX signals were visualized in MO-LAS-B cells treated with 25 nM paclitaxel (paclitaxel +). Untreated cells (paclitaxel -) are shown as a control. The arrowhead in the DAPI image represents potential CCF. **(c)** Percentages of CCF-positive cells treated or untread with paclitaxel. As shown in **panel b**, paclitaxel-treated and untread cells were subjected to IF experiments to visualize DAPI and γH2AX. **(d)** SA-β-gal signals in MO-LAS-B cells treated with 25 nM paclitaxel (paclitaxel +). SA-β-gal signals were visualized by SPiDER-βGal. Flow cytometric data analysis is shown in **Figure S3**. **(e)** Percentages of SA-β-gal-positive cells treated or untread with paclitaxel. SA-β-gal-positive cells were estimated, as shown in **Figure S3**. **(f)** Up-regulation of SASP genes in MO-LAS-B cells treated with 25 nM paclitaxel. Cells cultured in the presence and absence of paclitaxel were examined by RT-qPCR.

### ABT-263 vulnerability of angiosarcoma cells treated with cisplatin and paclitaxel

The above results indicate that angiosarcoma cells become senescent by cisplatin and paclitaxel treatment. We next explored senolytic agents to efficiently eliminate senescent angiosarcoma cells. To this end, we employed three recently identified senolytic agents, ABT-263 (inhibitor of anti-apoptotic BCL-2), BPTES (inhibitor of glutaminase 1), and RSL3 (inhibitor of GPX4 ferroptosis regulator) (Chang et al., 2016, Johmura et al., 2021, Liao et al., 2022, Zhu et al., 2016).

First, angiosarcoma cells were treated with either cisplatin or paclitaxel, followed by treatment with one of the three senolytic agents, as shown in the schematic procedure (**Figure 4a**). Drug-response curves indicate that MO-LAS-B and ISO-HAS-B cells treated with cisplatin or paclitaxel were more sensitive to ABT-263 than untreated control cells (**Figure 4b**); ABT-263 IC50 values were significantly lower for cisplatin- and paclitaxel-treated cells compared to untreated cells (**Figure 4c**). For instance, the ABT-263 IC50 values for cisplatin-treated MO-LAS-B cells and untreated cells were 1.58 and 14.0 μM, respectively, indicating that an 8.86-fold lower concentration of ABT-263 is sufficient to equivalently eliminate cisplatin-treated MO-LAS-B cells compared to untreated cells. To assess the efficacy of senolytic agents, a senolytic index is calculated by dividing the IC50 value of a particular senolytic agent for untreated control cells by that for drug-treated cells (Materials and Methods) (Jochems et al., 2021). The senolytic indexes of ABT-263 were 8.86 and 8.03 for cisplatin-treated MO-LAS-B and ISO-HAS-B cells, respectively, and 3.63 for paclitaxel-treated MO-LAS-B cells, indicating that at the conditions we tested cisplatin-treated angiosarcoma cells are more vulnerable to ABT-263 compared to paclitaxel-treated cells (**Figure 4c**).

**Figure 4.**
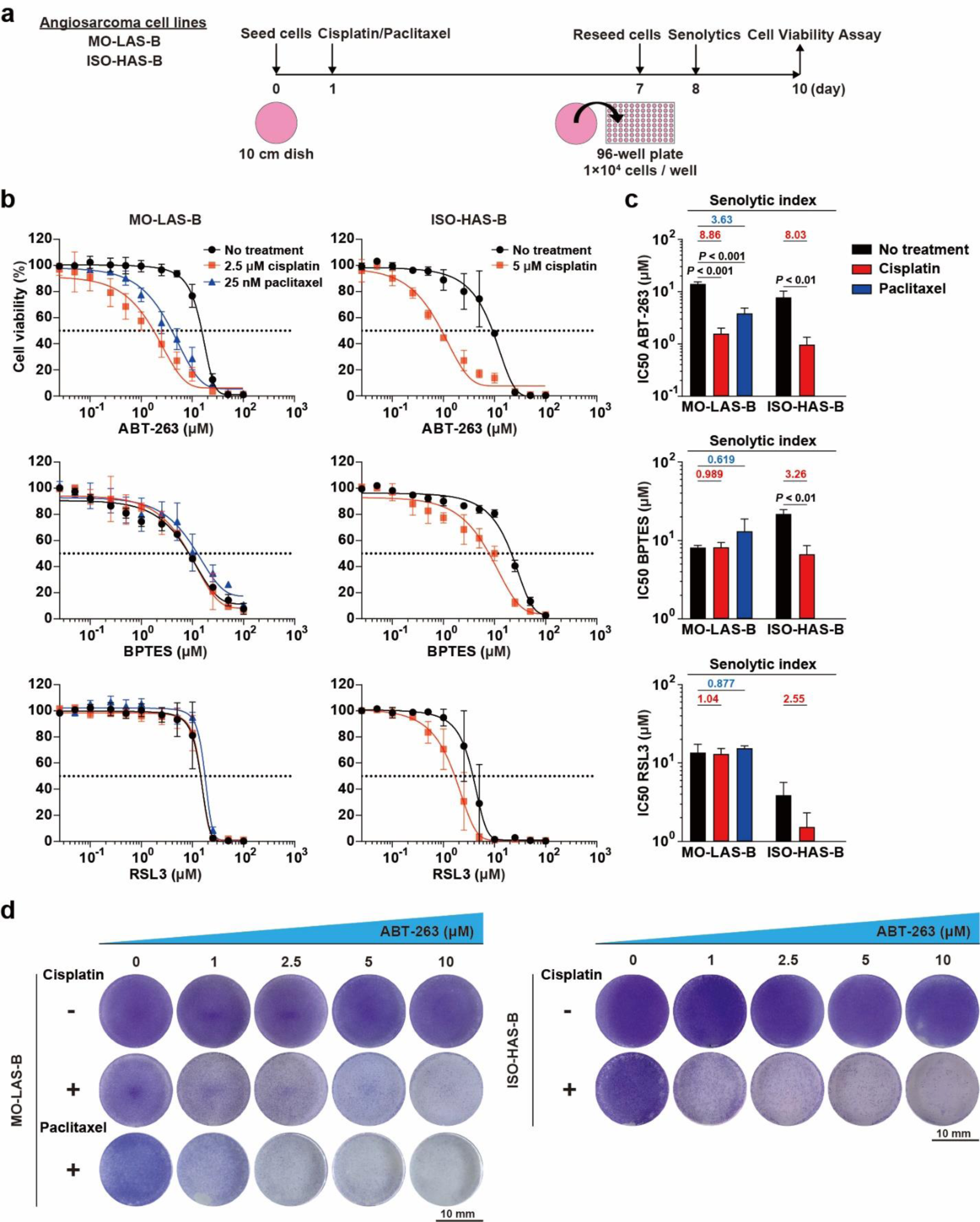
Cisplatin- and paclitaxel-treated angiosarcoma cells are vulnerable to ABT-263. **(a)** Schematic representation of the experimental procedures employed in **panels b-d** (see also Materials and Methods). **(b)** Dose-response curves of senolytic agents for MO-LAS-B and ISO-HAS-B cells. Cells treated with a chemotherapeutic drug, cisplatin or paclitaxel, were subsequently subjected to senolytic treatment either with ABT-263 (top), BPTES (middle), or RSL3 (bottom). Cells untreated with the chemotherapeutic drugs are also shown as a control. **(c)** IC50 values of the indicated senolytic agents for MO-LAS-B and ISO-HAS-B cells. The IC50 values were determined based on the dose-response curves in **panel b**. Numbers in red and blue indicate senolytic indexes for cisplatin- and paclitaxel-treated cells, respectively. Senolytic indexes were calculated as described in Materials and Methods. **(d)** Effect of ABT-263 treatment on colony formation of MO-LAS-B (left) and ISO-HAS-B cells (right) treated with cisplatin or paclitaxel. Quantification of the data is shown in **Figure S4**.

Moreover, the BPTES IC50 value for cisplatin-treated ISO-HAS-B cells was significantly lower than that for untreated control cells (senolytic index = 3.26), whereas BPTES had a negligible effect on cisplatin- and paclitaxel-treated MO-LAS-B cells (**Figure 4b-c**). Similarly, RSL3 also showed a moderate effect on cisplatin-treated ISO-HAS-B cells (senolytic index = 2.55), although the effect was not statistically significant (*p* = 0.10; **Figure 4b-c**). These results indicate that cisplatin and ABT-263 are the most effective combination for chemo-senolytic treatment of angiosarcoma cells among the combinations we tested.

Next, we examined the effect of ABT-263 on colony formation of angiosarcoma cells treated with cisplatin or paclitaxel and observed that colony formation of cisplatin- or paclitaxel-treated angiosarcoma cells was significantly impaired as ABT-263 concentration increased, although untreated control cells formed colonies regardless of increasing concentrations of ABT-263, clearly indicating that cisplatin- and paclitaxel-treatment renders angiosarcoma cells vulnerable to ABT-263 (**Figures 4d** and **S4**).

### ABT-263 eliminates cisplatin-induced senescent angiosarcoma cells via apoptosis

Based on the above results, we hypothesized that ABT-263 senolytic treatment would eliminate senescent angiosarcoma cells by inducing the apoptotic pathway. To test this hypothesis, cisplatin-treated cells and cells treated sequentially with cisplatin and ABT-263 were subjected to SA-β-gal staining. We observed that SA-β-gal-positive cells increased with cisplatin treatment, while they decreased with sequential treatment with cisplatin and ABT-263 (**Figure 5a**). To further investigate the effect of sequential treatment on angiosarcoma cells, we subjected the cells to DAPI and SA-β-gal staining and found that apoptotic cells with a punctate DAPI-staining pattern were readily detected after sequential treatment compared to cisplatin treatment alone (**Figure 5b**). Note that apoptotic cells with punctate DAPI signals were SA-β-gal-negative, indicating that ABT-263 treatment converts senescent cells to apoptotic cells, in which SA-β-gal signals are no longer detected (**Figure 5b**). Moreover, SA-β-gal-positive cells were significantly reduced from 72 to 23% for MO-LAS-B cells and from 79 to 28% for ISO-HAS-B cells, respectively, with sequential treatment compared with cisplatin treatment alone (**Figure 5c-d**).

**Figure 5.**
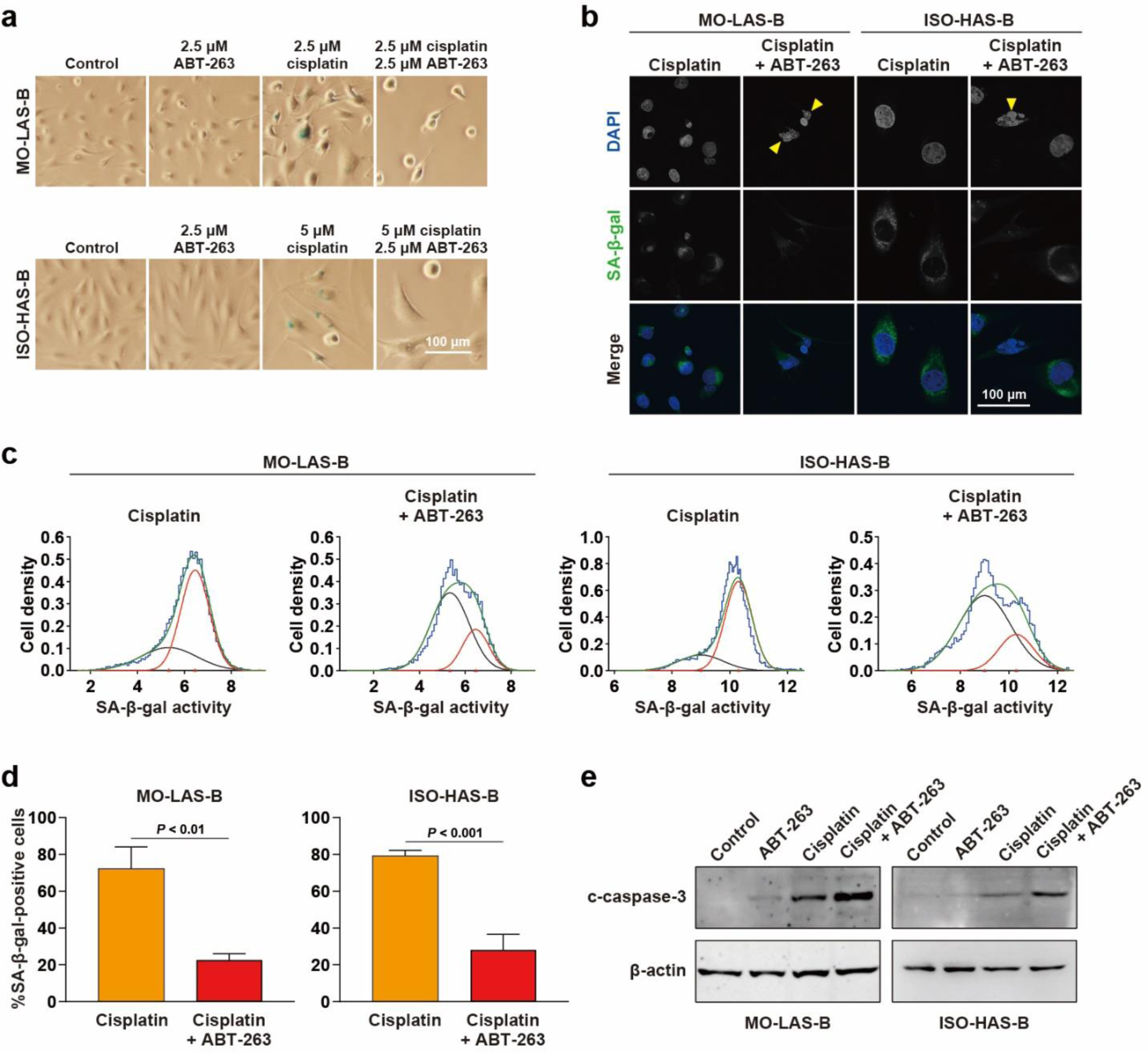
ABT-263 eliminates cisplatin-induced senescent angiosarcoma cells via apoptosis. **(a)** Effect of ABT-263 on angiosarcoma cells treated with cisplatin. MO-LAS-B (top) and ISO-HAS-B cells (bottom) were treated with ABT-263 or cisplatin alone or sequentially treated with cisplatin and ABT-263, as shown in Figure 4a, followed by SA-β-gal staining. Untreated cells were also subjected to the same analysis (control). **(b)** DAPI and SA-β-gal staining of cells treated with cisplatin alone or sequentially treated with cisplatin and ABT-263. SA-β-gal signals were visualized by SPiDER-βGal. Arrowheads represent apoptotic cells with a typical, punctate DAPI-staining pattern. **(c)** Reduction of a cisplatin-induced senescent cell population by ABT-263 treatment. MO-LAS-B and ISO-HAS-B cells treated with cisplatin or with the two agents (cisplatin and ABT-263) were subjected to SA-β-gal staining, followed by flow cytometric analysis. A ratio of the two populations, SA-β-gal-positive senescent and negative non-senescent cells, was estimated using the “mixdist” package in R. Briefly, flow cytometric data (blue) was used to estimate an entire cell population (green), which was in turn divided into two subset populations representing SA-β-gal-positive senescent (red) and negative non-senescent cells (black). **(d)** Percentages of SA-β-gal-positive cells treated or untreated with ABT-263 after cisplatin treatment. SA-β-gal-positive cells were estimated as described in **panel c**. **(e)** Western blot analysis monitoring an apoptosis marker, c-caspase-3, in angiosarcoma cells treated by the indicated agents. β-actin is a loading control.

In addition, we examined how different treatments promote apoptosis by following an apoptotic marker, c-capase-3, and observed that the apoptotic marker was most enhanced in cells sequentially treated with cisplatin and ABT-263 compared to other treatments, including cisplatin treatment alone (**Figure 5e**). This and the above results collectively suggest that cisplatin treatment converts angiosarcoma cells to senescent cells, which are in turn eliminated by ABT-263 senolytic treatment via promoting the apoptotic pathway. We also examined how angiosarcoma cells respond to treatment with varying concentrations of cisplatin and observed that the apoptotic marker was most enhanced at the 10 μM cisplatin condition, in which the punctate DAPI pattern was also detected, indicating that a higher dose of cisplatin is required for cisplatin single treatment to strongly promote the apoptotic pathway compared to sequential treatment (2.5 μM for MO-LAS-B cells and 5 μM for ISO-HAS-B cells; **Figure S5**).

### Paclitaxel-induced senescent angiosarcoma cells are eliminated by ABT-263 via apoptosis

We observed that SA-β-gal-positive cells increased by paclitaxel treatment but decreased by sequential treatment with paclitaxel and ABT-263 (**Figure 6a**). Moreover, apoptotic cells with a punctate DAPI-staining pattern were readily detected after sequential treatment (**Figure 6b**). In addition, SA-β-gal-positive cells were significantly reduced from 61 to 28% with sequential treatment compared with paclitaxel treatment alone (**Figure 6c-d**).

**Figure 6.**
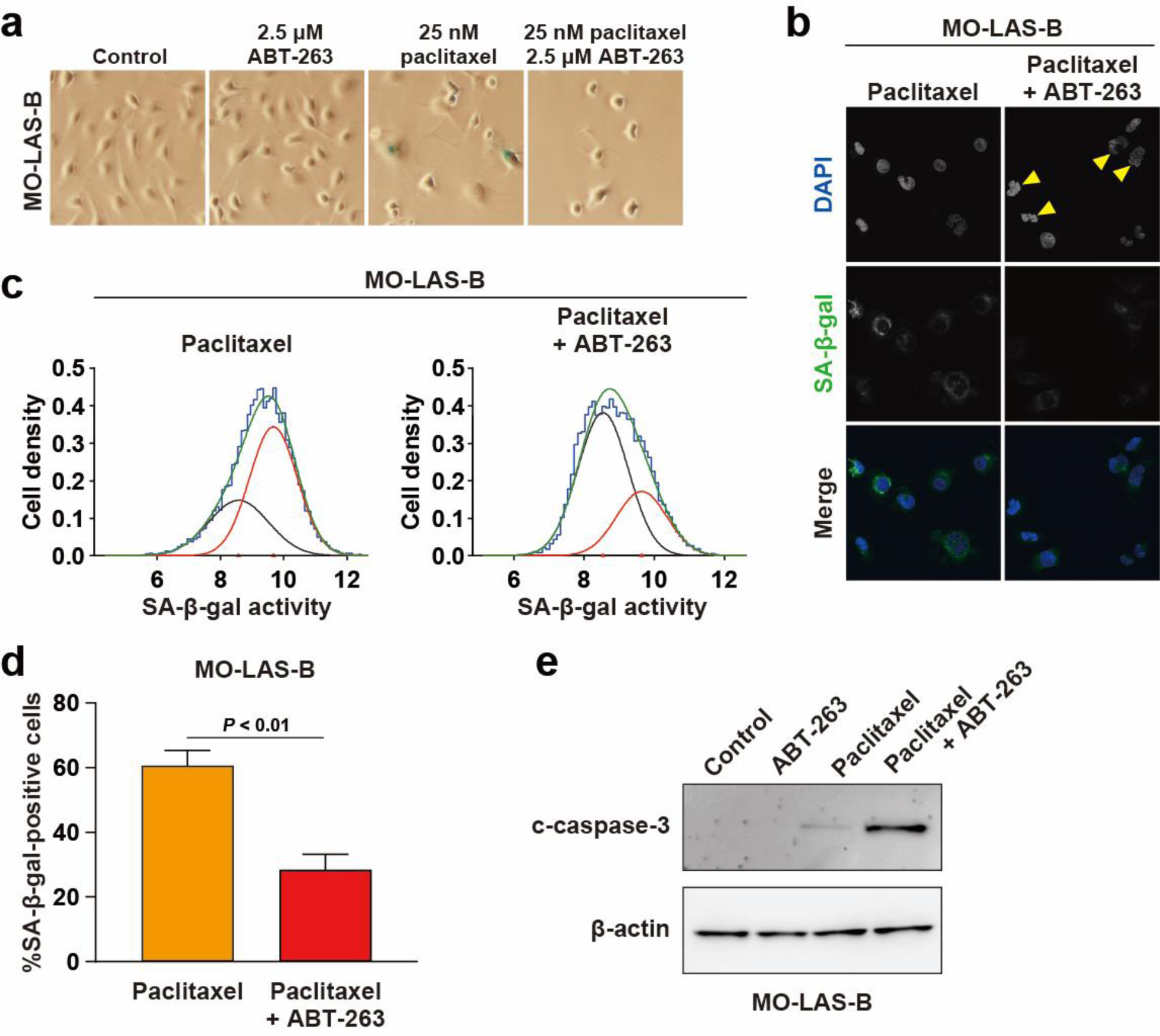
ABT-263 eliminates paclitaxel-induced senescent angiosarcoma cells via apoptosis. **(a)** Effect of ABT-263 on MO-LAS-B angiosarcoma cells treated with paclitaxel. The experiments were performed as described in Figure 4a. MO-LAS-B cells were treated with ABT-263 or paclitaxel alone or sequentially treated with paclitaxel and ABT-263, followed by SA-β-gal staining. **(b)** DAPI and SPiDER-βGal staining of MO-LAS-B cells treated with paclitaxel alone or sequentially treated with paclitaxel and ABT-263. Arrowheads represent apoptotic cells with a punctate DAPI-staining pattern. **(c)** Reduction of a paclitaxel-induced senescent cell population by ABT-263 treatment. The analysis was carried out as described in Figure 5c. **(d)** Percentages of SA-β-gal-positive senescent cells treated or untreated with ABT-263 after paclitaxel treatment. SA-β-gal-positive cells were estimated from the flow cytometric data in **panel c**. **(e)** Western blot analysis monitoring an apoptosis marker, c-caspase-3, in MO-LAS-B cells treated by the indicated agents.

We also examined the apoptotic marker and observed that c-capase-3 clearly increased in cells sequentially treated with paclitaxel and ABT-263 (**Figure 6e**). Again, our results indicate that ABT-263 eliminates senescent angiosarcoma cells induced by paclitaxel treatment via promoting the apoptotic pathway. In terms of paclitaxel single treatment, the apoptotic marker became much enhanced at paclitaxel concentrations of 50 and 100 nM, which are higher than the 25 nM paclitaxel condition used for sequential treatment (**Figure S6**), indicating that a higher dose of paclitaxel is required to induce apoptosis for paclitaxel treatment alone compared to sequential treatment.

### Suppression of interferon pathway genes by ABT-263 senolytic treatment

To further understand how treatment with chemotherapeutic and senolytic agents affects gene expression in angiosarcoma cells, we performed RNA-seq analysis. The degree of data point proximity on the principal component analysis (PCA) plot indicated that overall expression patterns were more similar within the same lines, irrespective of the treatment conditions (left panel in **Figure S7a**). The cell line-specific global expression patterns probably reflect the different origins of these angiosarcoma cell lines: MO-LAS-B and ISO-HAS-B cells are derived from lymphangiosarcoma and hemangiosarcoma, respectively (Masuzawa et al., 1999, Masuzawa et al., 2012). We also noticed that biological replicates were generally located in proximity, indicating the reproducibility of our RNA-seq analysis (middle and right panels in **Figure S7a**).

Although the overall expression patterns were more similar within the same angiosarcoma cell lines than those from the two different cell lines, it is possible that treatment with chemotherapeutic and senolytic agents has similar effects on some specific genes. To verify this possibility, we compared the expression profiles from the different treatment conditions within the same cell lines and identified differentially expressed genes (DEGs) based on a threshold of FDR < 0.05 and log2 (Fold-change) > 1.2 or < -1.2 (**Figure S7b**). Among DEGs, we first noticed that several SASP genes were up-regulated in angiosarcoma cells treated with cisplatin and paclitaxel (**Figure 7a**). This result indicates that chemotherapeutic drug treatment induces senescence of angiosarcoma cells and leads to the up-regulation of SASP genes, although activated SASP genes and the degrees of activation vary among the different conditions and the cell lines.

**Figure 7.**
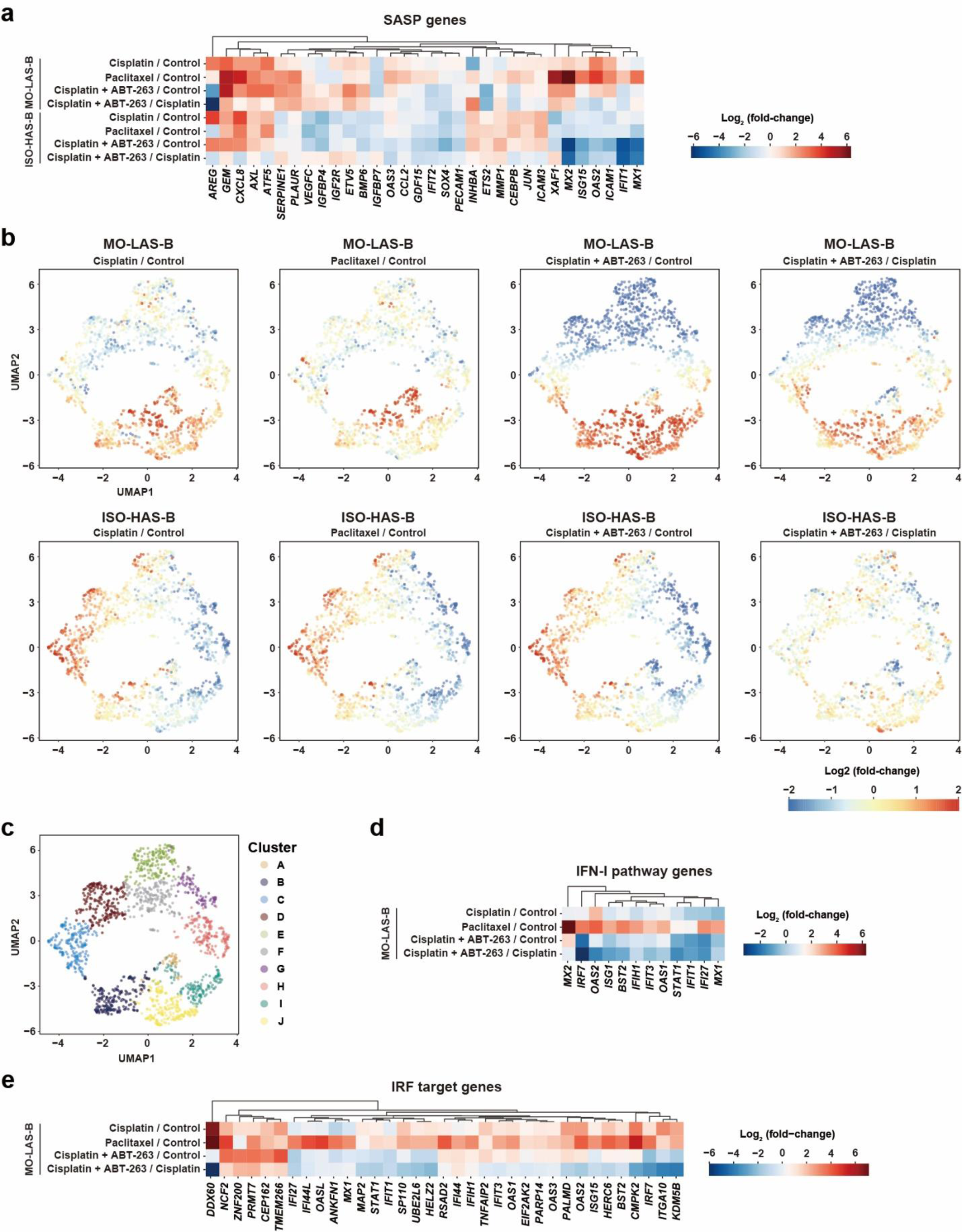
ABT-263 down-regulates genes in the IFN-I pathway. **(a)** Effect of chemotherapeutic drugs (cisplatin and paclitaxel) and senolytic agent (ABT-263) on the expression of SASP genes in angiosarcoma cells. MO-LAS-B and ISO-HAS-B cells were treated with the indicated agents and subjected to RNA-seq analysis. Expression data from untreated cells serves as a control. Expression fold changes of the SASP genes between the indicated conditions are shown. **(b)** Uniform Manifold Approximation and Projection (UMAP) plot of 1,662 differentially expressed genes (DEGs) based on log2 (fold-change scores). Each dot indicates one DEG. Genes with similar expression patterns across treatment conditions are positioned in proximity. **(c)** *K*-means clustering of DEGs. DEGs in Figure 7b were divided into 10 clusters based on the UMAP results. DEGs in each cluster are represented by the same color. **(d)** Effect of the indicated treatment on the expression of type-I interferon (IFN-I) pathway genes in MO-LAS-B cells. **(e)** Effect of the indicated treatment on the expression of interferon-regulatory factor (IRF) target genes.

To further explore how gene expression is affected by different treatments, we performed gene clustering analysis using the Uniform Manifold Approximation and Projection (UMAP) algorithm based on expression fold-changes of a total of 1,662 DEGs, which were defined by the comparison of RNA-seq data shown in **Figure S7b** (**Figure 7b**). In the UMAP plots, DEGs with similar expression changes were located nearby (**Figure 7b**). We found that up- and down-regulation of DEGs after treatment with cisplatin and paclitaxel were largely consistent within the same cell lines but were different between MO-LAS-B and ISO-HAS-B cells. This result suggests that the two cell lines differ in global gene expression responses to cisplatin and paclitaxel. Similarly, DEGs in cells treated with ABT-263 after chemotherapeutic drug treatment were largely different between the two cell lines.

We next sought to identify DEGs commonly affected by different treatments and cell lines. For this purpose, we performed *K*-means clustering, which divided DEGs into 10 clusters based on expression fold-changes caused by different treatments within the same cell lines (**Figure 7c**). Among the 10 clusters, cluster A was significantly enriched with SASP genes (7 SASP genes in 48 DEGs) compared to clusters B-J (25 SASP genes in 1,614 DEGs) (*p* < 0.001, chi-square test; **Figure S7c**). DEGs in cluster A were up-regulated in MO-LAS-B cells treated with cisplatin and down-regulated in the both cell lines after ABT-263 treatment (**Figure 7b**). To characterize DEGs in respective clusters, especially in cluster A, we performed gene ontology (GO) analysis and found that cluster A was enriched with genes associated with the type-I interferon (IFN-I) pathway (**Table 1**). Some DEGs in the IFN-I pathway showed similar expression responses to different treatments (**Figures 7d** and **S7d**). For instance, DEGs in the IFN-I pathway were up-regulated by chemotherapeutic drug treatment, especially with paclitaxel, and down-regulated by ABT-263 treatment. In addition, GO analysis of other clusters showed that genes in the cell-cycle- and DNA replication-regulation pathways were significantly enriched with clusters B, C, and D (**Table S1**). Since DEGs in cluster B, C, and D were up-regulated in ISO-HAS-B cells treated with chemotherapeutic drugs, the up-regulation of these genes are likely reflective of the proliferation inhibition caused by treatment (**Figure 7b**).

**Table 1.** GO analysis of genes in cluster A.

| Pathway | GO ID | Gene number | *p*-value |
| --- | --- | --- | --- |
| Defense response to virus | GO:0051607 | 18 | $7.26 \times 10^{-21}$ |
| Defense response to symbiont | GO:0140546 | 18 | $7.71 \times 10^{-21}$ |
| Response to virus | GO:0009615 | 19 | $4.76 \times 10^{-20}$ |
| Negative regulation of viral genome replication | GO:0045071 | 11 | $7.92 \times 10^{-19}$ |
| Negative regulation of viral process | GO:0048525 | 12 | $2.60 \times 10^{-18}$ |
| Regulation of viral genome replication | GO:0045069 | 11 | $1.23 \times 10^{-16}$ |
| Viral genome replication | GO:0019079 | 12 | $2.32 \times 10^{-16}$ |
| Regulation of viral process | GO:0050792 | 12 | $4.71 \times 10^{-15}$ |
| Regulation of viral life cycle | GO:1903900 | 11 | $3.95 \times 10^{-14}$ |
| Viral process | GO:0016032 | 15 | $5.19 \times 10^{-14}$ |
| Response to type I interferon | GO:0034340 | 9 | $4.30 \times 10^{-13}$ |
| Viral life cycle | GO:0019058 | 13 | $5.49 \times 10^{-13}$ |
| Cellular response to type I interferon | GO:0071357 | 8 | $1.36 \times 10^{-11}$ |
| Interferon-mediated signaling pathway | GO:0140888 | 8 | $1.15 \times 10^{-10}$ |
| Positive regulation of type I interferon production | GO:0032481 | 7 | $1.55 \times 10^{-10}$ |
| Positive regulation of interferon-beta production | GO:0032728 | 6 | $4.22 \times 10^{-10}$ |
| Type I interferon-mediated signaling pathway | GO:0060337 | 7 | $6.72 \times 10^{-10}$ |
| Cytosolic pattern recognition receptor signaling pathway | GO:0002753 | 7 | $3.28 \times 10^{-9}$ |
| Interferon-beta production | GO:0032608 | 6 | $4.80 \times 10^{-9}$ |
| Regulation of interferon-beta production | GO:0032648 | 6 | $4.80 \times 10^{-9}$ |
| Regulation of type I interferon production | GO:0032479 | 7 | $6.20 \times 10^{-9}$ |
| Type I interferon production | GO:0032606 | 7 | $6.20 \times 10^{-9}$ |
| MDA-5 signaling pathway | GO:0039530 | 4 | $1.06 \times 10^{-8}$ |
| Regulation of response to biotic stimulus | GO:0002831 | 10 | $1.50 \times 10^{-7}$ |
| Regulation of nuclease activity | GO:0032069 | 4 | $2.31 \times 10^{-7}$ |

Note that all the listed pathways are associated with IFN-I.

As shown above, cluster A was enriched with genes in the IFN-I pathway, and we also found that 35 genes in cluster A (total 48 genes) were targets of the interferon-regulatory factor (IRF) family of transcription factors (**Figure 7e**). It is known that the IRF transcription factors are involved in the induction of IFN-I pathway genes (McNab et al., 2015). Importantly, these IRF target genes were generally up-regulated by chemotherapeutic drug treatment and down-regulated by sequential treatment with cisplatin and ABT-263 compared to cisplatin treatment alone (**Figure 7e**). Since ABT-263 treatment eliminates senescent angiosarcoma cells, the observed down-regulation of IFN-I pathway genes after senolytic treatment is likely due to the elimination of cells expressing IFN-I pathway genes.

## Discussion

This study shows that angiosarcoma cells treated with a chemotherapeutic drug, cisplatin or paclitaxel, become senescent, and these senescent angiosarcoma cells are vulnerable to ABT-263 senolytic agent. The other senolytic agents, BPTES and RSL3, show a moderate senolytic effect on senescent angiosarcoma cells. Interestingly, senescent angiosarcoma cells induced by cisplatin exhibit more vulnerability to ABT-263 than those induced by paclitaxel, although paclitaxel is a commonly used chemotherapeutic drug for angiosarcoma treatment. Moreover, expression analysis predicts that senescent angiosarcoma cells expressing important senescence genes in the IFN-I pathway are eliminated by ABT-263 senolytic treatment.

Chemo-senolytic treatment has emerged as a potential therapy for cancer (Di Micco et al., 2021, Paez-Ribes et al., 2019, Wang et al., 2022). This combinational treatment consists of a one-two punch: chemotherapeutic drug treatment to induce senescence is followed by senolytic treatment to eliminate senescent cancer cells (Wang et al., 2022). It is well-accepted that genotoxic drug treatment of cancer cells induces senescence through a DNA damage response (Hernandez-Segura et al., 2018, Munoz-Espin and Serrano, 2014), and we observed that cisplatin-induced senescent angiosarcoma cells were more sensitive to ABT-263 senolytic agent than paclitaxel-induced senescent cells (**Figure 4**). This observed difference between cisplatin and paclitaxel can be explained by the efficiency of senescence induction. For instance, MO-LAS-B cells treated with 2.5 μM cisplatin and 25 nM paclitaxel showed 79 and 65% cell viabilities, respectively, implying that 25 nM paclitaxel is a slightly stronger condition than 2.5 μM cisplatin (**Figure 1b, e**). However, 67% and 48% of SA-β-gal-positive cells were detected after the 2.5 μM cisplatin and 25 nM paclitaxel treatment (**Figures 2e** and **3e**), suggesting that cisplatin is a more effective inducer for senescence of angiosarcoma cells than paclitaxel. Moreover, this difference in the efficiency of senescence induction may reflect the mode of action of these agents; cisplatin directly binds to DNA and induces DNA damage (Rottenberg et al., 2021), while paclitaxel inhibits mitotic spindle assembly and causes mitotic arrest (Jordan and Wilson, 2004). In any case, this study suggests that cisplatin is a better choice for a chemo-senolytic combinational approach than paclitaxel, at least for the conditions and the angiosarcoma cell lines we tested.

To eliminate senescent cells derived from chemotherapeutic drug treatment of angiosarcoma cells, we examined three senolytic agents: ABT-263 (Chang et al., 2016, Zhu et al., 2016), BPTES (Johmura et al., 2021), and RSL3 (Liao et al., 2022). We observed that all three agents could eliminate senescent angiosarcoma cells to varying degrees; ISO-HAS-B angiosarcoma cells treated by cisplatin exhibit enhanced vulnerability to the senolytic agents compared to untreated control cells, although ABT-263 was more effective than BPTES and RSL3 (**Figure 4b-c**). On the other hand, another angiosarcoma cell line (MO-LAS-B) treated with cisplatin or paclitaxel is only sensitive to ABT-263. This broader spectrum of ABT-263 is probably because the inhibition of anti-apoptotic Bcl-2 family proteins has a wider range for the elimination of senescent cells. In this regard, it has been shown that senescent cells become resistant to apoptosis via the up-regulation of Bcl-2 family proteins (Ryu et al., 2007). Taken together, cisplatin and ABT-263 are the best combination among the treatments we tested for angiosarcoma cells.

Expression analysis shows that SASP genes are up-regulated by chemotherapeutic drug treatment and that genes in the IFN-I pathway are consistently down-regulated in the angiosarcoma cell lines after ABT-263 treatment. This down-regulation of IFN-I pathway genes is explained by the elimination of senescent cells expressing IFN-I pathway genes by ABT-263 senolytic treatment. Since the IFN-I pathway is activated in senescent cells and plays a pivotal role in the expression of SASP factors, the elimination of senescent cells expressing IFN-I pathway genes can contribute to the suppression of the SASP and thus have a positive impact on the tumor microenvironment (De Cecco et al., 2019). For instance, some SASP factors are known to promote immune evasion; IL-6 secreted by senescent stroma cells contributes to the recruitment of suppressive myeloid cells to the tumor microenvironment (TME) and compromises the anti-tumor immune surveillance (Ruhland et al., 2016).

Our study shows that treatment with chemotherapeutic drugs (cisplatin and paclitaxel) causes senescence of angiosarcoma cells and that ABT-263 senolytic treatment allows for the efficient removal of senescent angiosarcoma cells. Therefore, a combinational approach is promising for angiosarcoma treatment. Importantly, a lower dose of cisplatin is sufficient to induce senescence compared to the dose to elicit apoptosis and should ease deleterious side effects from the treatment. Although ABT-263 has platelet toxicity, recent advances have been made to reduce the ABT-263 toxicity (Gonzalez-Gualda et al., 2020, Munoz-Espin et al., 2018). In the future, many more senolytic agents will be developed, and our next challenge is to find the optimal combination with the least toxicity to efficiently eliminate angiosarcoma cells using the combinational approach.

## Materials and Methods

### Cell lines and culture conditions

The MO-LAS-B lymphangiosarcoma cell line and ISO-HAS-B hemangiosarcoma cell line were obtained from the Cell Resource Center for Biomedical Research, Tohoku University. These cell lines were cultured in DMEM (Wako) supplemented with 10% FBS (Corning) and 1% penicillin-streptomycin mixed solution (Nacalai Tesque, #26252-94) at 37℃ in a humidified atmosphere containing 5% CO2.

### Cell viability

MO-LAS-B and ISO-HAS-B cells were first seeded in 96-well cell culture plates with DMEM at a density of 10,000 cells per well and cultured for 1 day to allow the cells to adhere to the culture plates. The cells were sequentially cultured for 48 hours in DMEM supplemented with one of the following chemicals: chemotherapeutic drugs [cisplatin (Selleck, S1166) and paclitaxel (Selleck, S1150)] and senolytic agents [ABT-263 (Selleck, S1001), BPTES (Selleck, S7753), and RSL3 (Selleck, S8155)], at the indicated concentrations. Cell numbers were measured using Synergy LX Multimode Reader (Agilent) and Cell Count Reagent SF Kit (Nacalai Tesque). IC50 values of the respective agents were estimated from three independent experiments using a nonlinear regression model.

For combinational treatment with two selected agents, MO-LAS-B cells were first cultured in 10 cm dishes with DMEM for 1 day to attach the cells to the culture dishes and subsequently cultured for 6 days in DMEM supplemented with the chemotherapeutic drugs, 2.5 μM cisplatin or 25 nM paclitaxel. ISO-HAS-B cells were cultured in a similar manner, except that medium containing 5 μM cisplatin was used. The medium containing each chemotherapeutic drug was exchanged every 3–4 days. The MO-LAS-B and ISO-HAS-B cells were next reseeded in 96-well cell culture plates at a density of 10,000 cells per well and cultured for 1 day to attach the cells to the culture plates. The cells were subsequently cultured in DMEM containing one of the senolytic agents (ABT-263, BPTES, or RSL3) at the indicated concentrations for 48 hours. Cell numbers and IC50 values were estimated as described above for single treatment. A senolytic index was calculated as the IC50 value of a senolytic agent without chemotherapeutic drug treatment divided by that with the treatment.

### Colony formation assay

MO-LAS-B and ISO-HAS-B cells were first seeded in 24-well cell culture plates with DMEM at a density of 5,000 cells per well and cultured for 1 day to attach the cells to the culture plates. They were sequentially cultured for 6 days in DMEM supplemented with one of the following chemicals: chemotherapeutic drugs (cisplatin and paclitaxel) and senolytic agent (ABT-263). The media containing respective agents at the indicated concentrations were exchanged every 3–4 days. Colonies were next washed twice with PBS and fixed with 4% paraformaldehyde (w/v) at room temperature for 15 minutes. The colonies were washed once with PBS and stained with 0.5% crystal violet at room temperature for 20 minutes. Colony formation was quantified as integrated intensity using NIH ImageJ software.

For combinational treatment with two selected agents, MO-LAS-B cells were first cultured in 10 cm dishes with DMEM for 1 day to attach the cells to the dishes and sequentially cultured for 6 days in DMEM supplemented with a chemotherapeutic drug, either 2.5 μM cisplatin or 25 nM paclitaxel. ISO-HAS-B cells were cultured in a similar manner, except that a medium with 5 μM cisplatin was used. The media containing the respective chemotherapeutic drugs were exchanged every 3–4 days. Subsequently, the MO-LAS-B and ISO-HAS-B cells were reseeded in 24-well cell culture plates at a density of 30,000 cells per well and cultured in DMEM for 1 day to attach the cells to the culture plates. The cells were next cultured in DMEM containing ABT-263 at the indicated concentrations for 4–6 days. The culture period was decided based on cell confluency. The media with ABT-263 were also exchanged every 3–4 days. Colony formation was examined as described above for single treatment.

### Senescence-associated (SA)-β-gal staining

SA-β-gal staining was performed as previously described (Dimri et al., 1995). Cells were seeded in 24-well cell culture plates with cover glasses (Marienfeld Superior, #0111520) and cultured under the indicated conditions. The cells were washed twice with PBS and fixed with a fixation solution containing 2% formaldehyde and 0.2% glutaraldehyde in PBS at room temperature for 5 minutes. The cells were next washed twice with PBS and stained with an X-gal staining solution [150nM NaCl, 2mM MgCl2, 5mM K3Fe(CN)6, 5mM K4Fe(CN)6, 40mM Na2HPO4 pH 6.0, and 1mg/mL X-gal (Sigma)] at 37℃ overnight in a non-CO2 incubator (EYELA, SLI-700).

### SA-β-gal quantification

SA-β-gal positive cells in cell populations were counted using Cellular Senescence Detection Kit SPiDER-βGal (Dojindo) (Doura et al., 2016). In brief, live cells were washed once with PBS and incubated with a Bafilomycin A1 working solution for 1 hour in a 5% CO2 incubator. The cells were next incubated with a SPiDER-βGal working solution for 30 minutes in the 5% CO2 incubator. After removing the supernatant, the cells were washed twice with PBS and harvested by trypsin treatment. The cells were resuspended with DMEM and analyzed on BD FACSCanto™ II Flow Cytometry System (BD Biosciences).

### Immunofluorescence

Cells were seeded on cover glasses (Marienfeld Superior, #0111520) and fixed with a 1:1 mixture of cold methanol and acetone at -20℃ for 5 minutes. The cells were washed three times with PBS at room temperature for 5 minutes and immersed in a PBS solution containing 1% BSA at 37℃ for 30 minutes. The cells were next incubated with a primary antibody, 1:500-diluted mouse anti-phospho-Histone H2A.X (Ser139) clone JBW301 (Sigma-Aldrich, #05-636), in a PBS solution containing 1% BSA at 37℃ for 1 hour. After the cells were washed three times with PBS for 5 minutes, they were incubated with a secondary antibody, 1:500-diluted Alexa Fluor 488-conjugated goat anti-mouse IgG H&L (Abcam, ab150113), in a PBS solution containing 1% BSA at 37℃ for 30 minutes. The cells were washed three times with PBS for 5 minutes and mounted on slide glasses (ASLAB, #1-9646-11) using Antifade Mounting Medium with DAPI (VECTASHIELD, H-1200). Immunofluorescence images were captured using a confocal microscope (Olympus, FV1000) with a 60x lens.

### Western blotting

Cells were suspended in PBS and sonicated using Bioruptor (Cosmo Bio, UCD-250HSA) at 4℃ for 30 minutes, repeating 30-second ON/OFF cycles. After centrifugation at 13,000 rpm (10,000 x g) for 30 minutes, the protein amount of the supernatant was quantified using Pierce™ BCA Protein Assay Kit (Thermo Fisher Scientific) and Synergy LX Multimode Reader (Agilent). Equal volumes of the supernatant, 3× protein sample buffer, and PBS were mixed to have a protein extract in 1× sample buffer (50 mM Tris-HCl pH 6.8, 15% sucrose, 2 mM EDTA pH 7.4, 3% SDS, 0.01% bromophenol blue, and 10% β-mercaptoethanol). The protein sample was heated at 95℃ for 2 minutes. Fifty micrograms of the respective samples were separated on 13% SDS-PAGE gel and transferred to an Immobilon®-P PVDF (polyvinylidene difluoride) membrane (Merk Millipore, IPVH00010) using XCell II™ Blot Module (Thermo Fisher Scientific) at 60 V for 120 minutes. The membrane was next treated with a PBS solution containing 5% non-fat milk at room temperature for 1 hour and was incubated with primary antibodies at room temperature overnight. The primary antibodies used in this study were 1:2,500-diluted mouse anti-β-actin (Wako, #017-24551) and 1:1,000-diluted rabbit anti-cleaved caspase-3 (Cell Signaling, #9661). The membrane was washed three times with PBS for 10 minutes and incubated with secondary antibodies at room temperature for 1 hour. The secondary antibodies used in this study were 1:2,500-diluted HRP-conjugated anti-mouse IgG (Promega, #W4021) and anti-rabbit IgG (Promega, #W4011). The membrane was washed three times with PBS for 10 minutes and subjected to signal detection using ECL™ Prime Western Blotting System (Cytiva) and ImageQuant™ LAS 4000 mini (GE HealthCare). The primary and secondary antibodies were diluted with a PBS solution containing 1% non-fat milk.

### RT-qPCR

Total RNA was extracted using PureLink™ RNA Mini Kit (Thermo Fisher Scientific) and subjected to reverse transcription using SuperScript™ II Reverse Transcriptase (Thermo Fisher Scientific) according to the manufacturer’s instructions. The cDNA was used as a template for quantitative PCR using KAPA SYBR FAST qPCR Kits (KAPA) and 7300 Real-Time PCR System (Applied Biosystems). Expression levels of target genes were normalized using an mRNA level of the β2-microglobulin gene (*B2M*) as an internal control. The 2^-ΔΔCT^ method was used to calculate the fold change of target genes (Livak and Schmittgen, 2001).

The sequences of RT-qPCR primers for the respective genes were as follows: *B2M* forward (5’-GGCATTCCTGAAGCTGACA-3’) and reverse (5’-CTTCAATGTCGGATGGATGAAAC-3’); *IL1B* forward (5’-AGCTCGCCAGTGAAATGATGG-3’) and reverse (5’- GTCCTGGAAGGAGCACTTCAT-3’); *IL6* forward (5’-ACATCCTCGACGGCATCTCA-3’) and reverse (5’-TCACCAGGCAAGTCTCCTCA-3’); *MMP1* forward (5’- AATAGTGGCCCAGTGGTTGA-3’) and reverse (5’-GGCTGCTTCATCACCTTCAG-3’); *CCL20* forward (5’-CTCCTGGCTGCTTTGATGTC-3’) and reverse (5’- TGCTTGCTGCTTCTGATTCG-3’); *OAS2* forward (5’-TTCTCCAGCCCAACAAATGC-3’) and reverse (5’-AGTCTTCAGAGCTGTGCCTT-3’); *OAS3* forward (5’- ACTGGAGTCGAGACCTAGGT-3’) and reverse (5’-TAGACCCACTCCCTCATCCA-3’); *ISG15* forward (5’-ACAAATGCGACGAACCTCTG-3’) and reverse (5’- CGCAGATTCATGAACACGGT-3’); *IFIT1* forward (5’-GCGGTTTCCACATGACAACT-3’) and reverse (5’-ATTCATGAGGGGCAGTCACA-3’); *MX1* forward (5’- CGGAATCTTGACGAAGCCTG-3’) and reverse (5’-CCTTTCCTTCCTCCAGCAGA-3’).

### RNA-seq

Total RNA was extracted using PureLink™ RNA Mini Kit (Thermo Fisher Scientific) according to the manufacturer’s instructions. The total RNA was treated with RQ1 RNase-Free DNase (Promega, #M6101) at 37°C for 40 minutes and purified by phenol/chloroform extraction and ethanol precipitation. 3’mRNA-seq libraries were prepared using QIAseq UPX 3’ Transcriptome Kit (QIAGEN). In brief, the purified total RNA was reverse-transcribed using an oligo-dT primer containing a sample barcode index and a unique molecule identifying sequence. The indexed cDNA samples prepared from different cell lines and culture conditions were combined, followed by adapter ligation and library amplification. The prepared libraries were sequenced on Illumina NextSeq2000 platform using a paired-end mode with a read length of 100 bp for transcripts and 27 bp for indexes, according to the library preparation kit instructions.

Sequenced data from respective samples were adjusted to 4.5 million reads by randomly selecting reads to minimize biases derived from variation in read numbers. Sequenced reads were aligned to the human reference genome GRCh38 using the STAR aligner (version 2.7.10a) (Dobin et al., 2013). The UMI Tools package (version 1.1.4) was used to demultiplex the pooled samples and remove redundant reads reflecting a potential PCR bias (Smith et al., 2017). Reads aligned to multiple positions were discarded using the SAMtools (version 1.14) (Li et al., 2009). Aligned reads were counted using the htseq-count script (version 2.0.3) based on the GENCODE v44 gene annotation (Anders et al., 2015, Frankish et al., 2021). The DESeq2 program (version 1.28.1) was employed to identify differentially expressed genes (Love et al., 2014).

### Gene ontology (GO)

GO analysis was performed to investigate how treatment with chemotherapeutic and senolytic agents affects genome-wide gene expression in angiosarcoma cells. RNA-seq data derived from different cell lines and culture conditions were subjected to the following analysis. First, RNA-seq data from the same cell lines were compared between cisplatin-treated and untreated control cells (cisplatin/control), paclitaxel-treated and control cells (paclitaxel/control), cisplatin- and ABT-263-treated and control cells (cisplatin+ABT-263/control), and cisplatin- and ABT-263-treated and cisplatin-treated cells (cisplatin+ABT-263/cisplatin). Differentially expressed genes (DEGs) were defined based on a threshold of FDR < 0.05 and log2 (Fold-change) > 1.2 or < -1.2 (red dots in **Figure S7b**). Second, the union of differentially expressed genes (n =1,662) in the MO-LAS-B and ISO-HAS-B cell lines was subjected to the UMAP algorithm to place DEGs at 2D coordinates, allowing DEGs with similar expression changes to be positioned nearby (McInnes et al., 2018). *K*-means clustering was performed on the UMAP result using the scikit-learn package (Pedregosa et al., 2011). Parameters, random_state and n_clusters, were set as 100 and 10, respectively. Gene ontology enrichment analysis was performed using the clusterProfiler package (Wu et al., 2021).

## Abbreviations

SASP: senescence-associated secretory phenotype
IFN-I: type-I interferon
NCCN: National Comprehensive Cancer Network
IC50: half maximal inhibitory concentration
GDSC: Genomics of Drug Sensitivity in Cancer
SA-β-gal: senescence-associated β-galactosidase
CCF: cytoplasmic chromatin fragments
PCA: principal component analysis
DEGs: differentially expressed genes
UMAP: Uniform Manifold Approximation and Projection
GO: gene ontology
IRF: interferon-regulatory factor

## Data availability

All relevant data that support the finding of this study are available from the authors upon request.

## Acknowledgments

We would like to thank Michiko Kanbayashi and Makiko Fukuuchi for their generous support in the laboratory. This work was supported by JSPS KAKENHI grant JP20K23376, Takeda Science Foundation, The Mitsubishi Foundation, and The Photo-excitonix Project in Hokkaido University, and The Grant for Joint Research Program of the Institute for Genetic Medicine, Hokkaido University (to K.N.). This work was also supported by JST SPRING grant JPMJSP2119 (to X.W.).

## Author contributions

X.W., T.F., and K.N. designed the experiments of this study. X.W., C.Y.C., A.Y., S.H., H.J., and S.O. performed the experiments. X.W., C.Y.C., H.T., and S.O. analyzed data. All authors contributed to writing and editing the manuscript.

## Competing interests

The authors declare no competing interests.

## Notes

### Competing Interest Statement

The authors have declared no competing interest.

