## Supplementary material for "Chemo-senolytic therapeutic potential against angiosarcoma": Figure S

### Supplemental Figures and Table

### Supplemental Figure 1

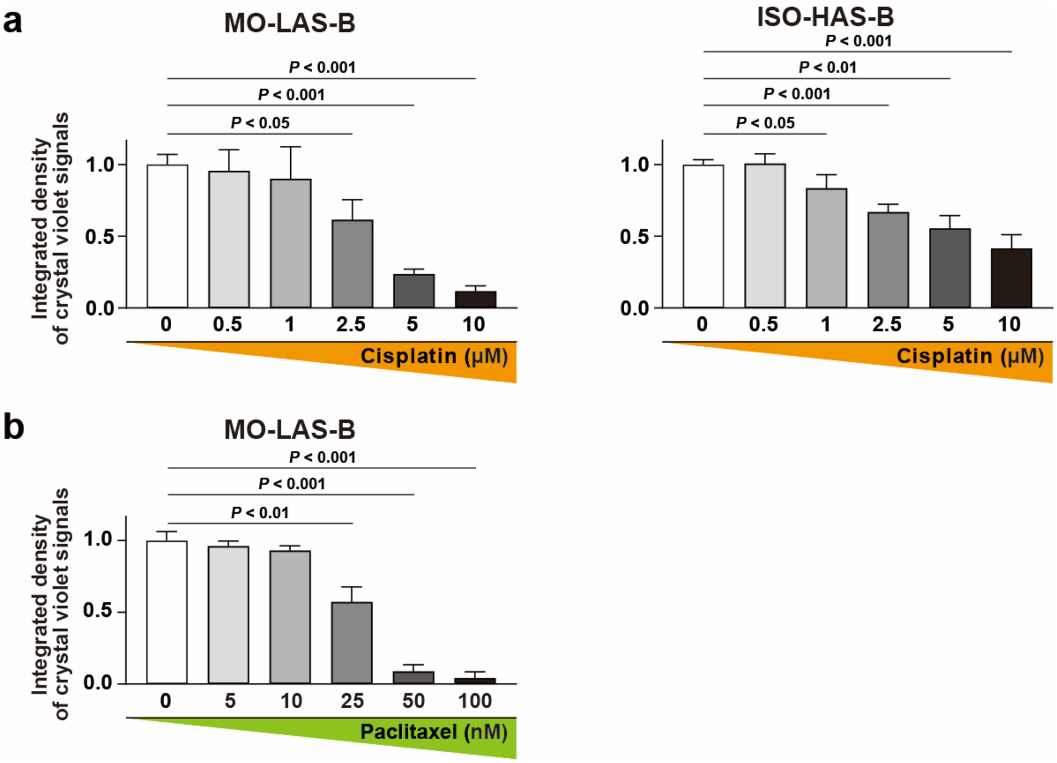


**Figure S1. Cisplatin and paclitaxel inhibit colony formation of angiosarcoma cells.**
**(a)** Effect of cisplatin treatment on colony formation. The experiments were performed as shown in **Figure 1a** (see Materials and Methods), and results from three biological replicates are shown. **(b)** Effect of paclitaxel treatment on colony formation. The experiments were performed as shown in **Figure 1d**.

**Supplemental Figure 2**

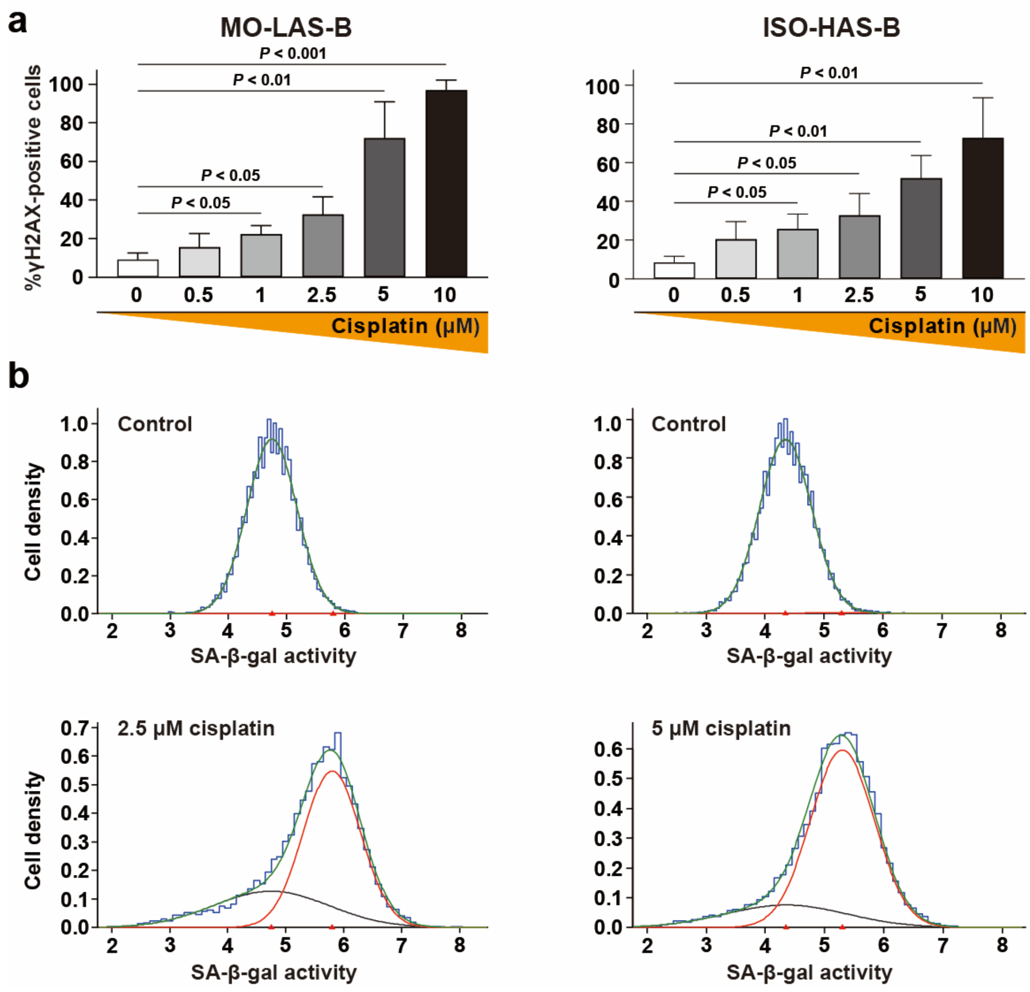


**Figure S2. Cisplatin-induced senescence was quantified by flow cytometry.**

**(a)** Percentages of γH2AX-positive cells after cisplatin treatment. The experiments were performed as shown in **Figure 2a**.

**(b)** Increase in SA-β-gal-positive cells after cisplatin treatment. MO-LAS-B (left) and ISO-HAS-B cells (right) were treated with 2.5 and 5 μM cisplatin, respectively (bottom), and subjected to SPiDER-βGal staining, followed by flow cytometric analysis. Untread control cells were also subjected to the same analysis (top). A ratio of the two populations, SA-β-gal-positive senescent and negative non-senescent cells, was estimated using the "mixdist" package in R. In brief, flow cytometric data (blue) was used to estimate an overall cell population (green), which was in turn divided into two subset populations reflecting SA-β-gal-positive senescent (red) and negative non-senescent cells (black). Percentages of senescent cell populations from three biological replicates are summarized in **Figure 2e**.

**Supplemental Figure 3**

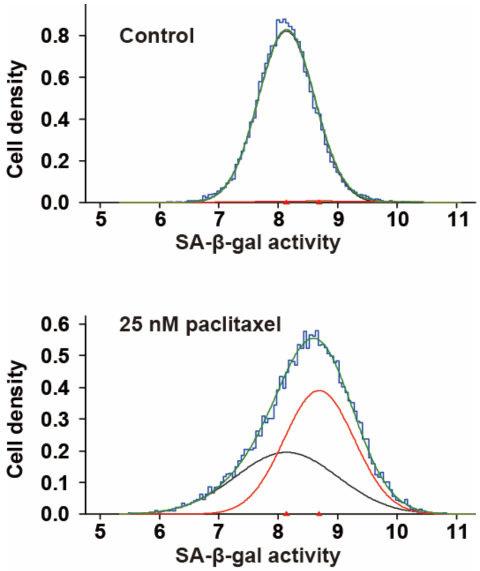


**Figure S3. Paclitaxel-induced senescence was quantified by flow cytometry.**

MO-LAS-B cells treated with 25 nM paclitaxel were subjected to SPiDER-βGal staining, followed by flow cytometric analysis. Untreated cells were also subjected to the same analysis (control). The analysis was carried out as described in **Figure S2b**, and results from three biological replicates are shown in **Figure 3e**.

**Supplemental Figure 4**

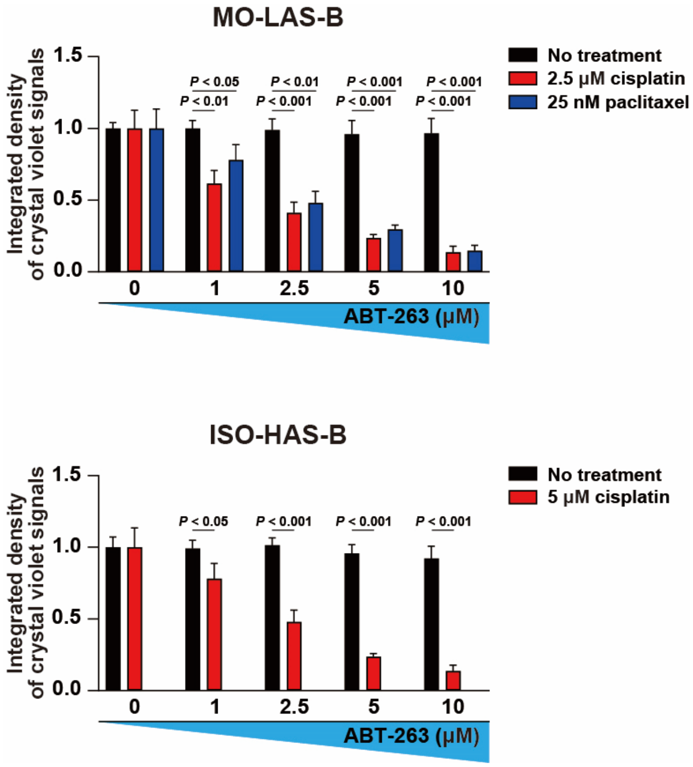


**Figure S4. ABT-263 inhibits colony formation of cisplatin- and paclitaxel-treated angiosarcoma cells.**

The experiments were performed as shown in **Figure 4d**, and results from three biological replicates are shown.

**Supplemental Figure 5**

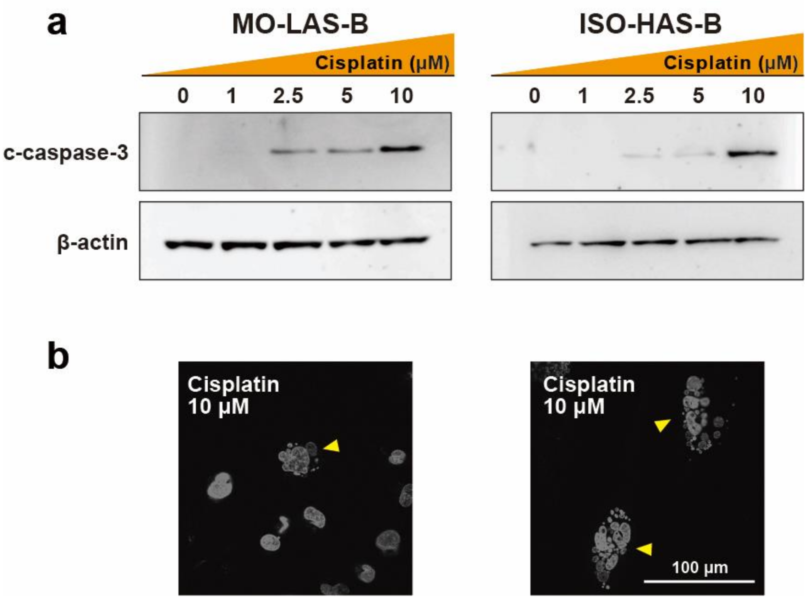


**Figure S5. A high dose of cisplatin induces apoptosis of angiosarcoma cells.**

**(a)** Monitoring an apoptosis marker, c-caspase-3, in angiosarcoma cells treated with cisplatin. MO-LAS-B and ISO-HAS-B cells were treated with cisplatin at the indicated concentrations for 6 days. The expression level of c-caspase-3 was examined by western blotting. β-actin is a loading control.

**(b)** Apoptotic cells detected after cisplatin treatment. MO-LAS-B (left) and ISO-HAS-B cells (right) were stained by DAPI after 10 μM cisplatin treatment. Arrowheads represent apoptotic cells with a typical, punctate DAPI-staining pattern.

**Supplemental Figure 6**

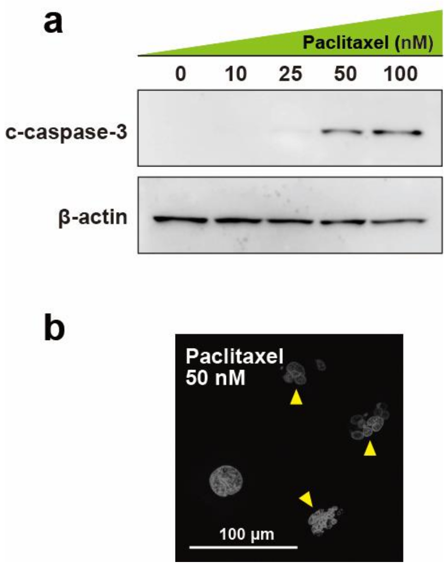


**Figure S6. A high dose of paclitaxel induces apoptosis of angiosarcoma cells.**

**(a)** Monitoring an apoptosis marker, c-caspase-3, in angiosarcoma cells treated by paclitaxel. MO-LAS-B cells were treated with paclitaxel at the indicated concentrations for 6 days. The expression level of c-caspase-3 was examined by western blotting.

**(b)** Apoptotic cells detected after paclitaxel treatment. MO-LAS-B cells were stained by DAPI after 50 nM paclitaxel treatment. Arrowheads represent apoptotic cells with a punctate DAPI-staining pattern.

**Supplemental Figure 7**

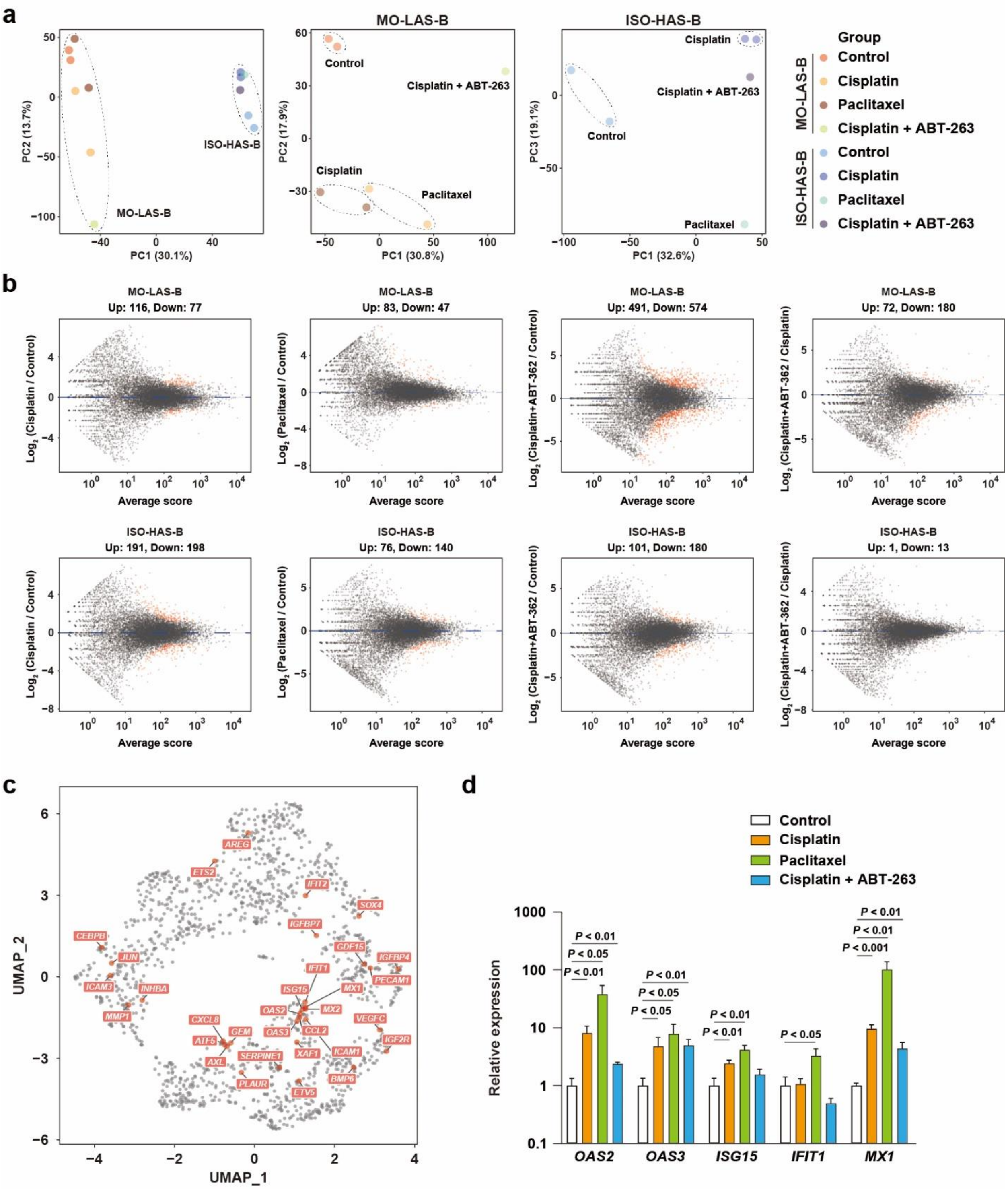


**Figure S7. Chemotherapeutic and senolytic agents affect gene expression in angiosarcoma cells.**

**(a)** Principal component analysis (PCA) of RNA-seq data. RNA-seq samples from the same cell lines were circled by dotted lines (left). In addition, RNA-seq samples from MO-LAS-B (middle) and ISO-HAS-B cells (right) were plotted separately, and biological replicates were circled. Data points in proximity indicate higher similarities among expression profiles of the relevant RNA-seq samples.

**(b)** $Log_2$ (expression fold-change) between the indicated conditions (Y-axis) were plotted against average expression levels of the respective genes (X-axis). Differentially expressed genes (DEGs; red) were defined based on the criteria of FDR < 0.05 and $log_2$ (fold-change) > 1.2 or < -1.2.

**(c)** SASP genes in the UMAP plot shown in **Figure 7b** and **c**.

**(d)** Effect of the indicated treatment on the expression of type-I interferon (IFN-I) pathway genes. These INF-I pathway genes also belong to SASP genes. RNA levels of the indicated genes in MO-LAS-B cells were quantified by RT-qPCR.

**Supplemental Table 1. GO analysis of cluster B-J genes**

| Pathway | GO ID | Gene number | *p*-value |
|---|---|---|---|
| **Cluster B** | | | |
| *DNA-templated DNA replication | GO:0006261 | 15 | $6.09\times10^{-11}$ |
| *DNA replication | GO:0006260 | 17 | $2.51\times10^{-9}$ |
| rRNA metabolic process | GO:0016072 | 16 | $5.84\times10^{-9}$ |
| *Mitotic cell cycle checkpoint signaling | GO:0007093 | 11 | $1.46\times10^{-7}$ |
| rRNA processing | GO:0006364 | 13 | $2.70\times10^{-7}$ |
| Ribosome biogenesis | GO:0042254 | 15 | $2.89\times10^{-7}$ |
| *Negative regulation of cell cycle G2/M phase transition | GO:1902750 | 8 | $3.10\times10^{-7}$ |
| *Negative regulation of cell cycle process | GO:0010948 | 15 | $6.33\times10^{-7}$ |
| *Mitotic G2/M transition checkpoint | GO:0044818 | 7 | $6.60\times10^{-7}$ |
| *DNA-templated DNA replication maintenance of fidelity | GO:0045005 | 7 | $1.27\times10^{-6}$ |
| **Cluster C** | | | |
| *Mitotic nuclear division | GO:0140014 | 22 | $1.91\times10^{-13}$ |
| *Nuclear division | GO:0000280 | 26 | $8.60\times10^{-13}$ |
| *Organelle fission | GO:0048285 | 27 | $1.28\times10^{-12}$ |
| *DNA replication | GO:0006260 | 20 | $8.71\times10^{-12}$ |
| *Negative regulation of chromosome organization | GO:2001251 | 13 | $1.10\times10^{-11}$ |
| *Sister chromatid segregation | GO:0000819 | 19 | $1.14\times10^{-11}$ |
| *Chromosome segregation | GO:0007059 | 24 | $1.84\times10^{-11}$ |
| *Negative regulation of sister chromatid segregation | GO:0033046 | 10 | $4.52\times10^{-11}$ |
| *Negative regulation of mitotic sister chromatid segregation | GO:0033048 | 10 | $4.52\times10^{-11}$ |
| *Negative regulation of mitotic metaphase/anaphase transition | GO:0045841 | 10 | $4.52\times10^{-11}$ |
| **Cluster D** | | | |
| Cytoplasmic translation | GO:0002181 | 10 | $3.43\times10^{-6}$ |
| Cellular detoxification | GO:1990748 | 7 | $1.28\times10^{-4}$ |
| Cellular response to toxic substance | GO:0097237 | 7 | $2.05\times10^{-4}$ |
| *DNA strand elongation involved in DNA replication | GO:0006271 | 3 | $3.71\times10^{-4}$ |
| Cellular oxidant detoxification | GO:0098869 | 6 | $4.01\times10^{-4}$ |
| Fibroblast proliferation | GO:0048144 | 6 | $4.95\times10^{-4}$ |

(*continued on next page*)

**Supplemental Table 1. GO analysis of cluster B-J genes (*continued*)**

| Pathway | GO ID | Gene number | *p*-value |
|---|---|---|---|
| Detoxification | GO:0098754 | 7 | $5.08{\times}10^{-4}$ |
| Oxidative phosphorylation | GO:0006119 | 7 | $5.99{\times}10^{-4}$ |
| Cellular respiration | GO:0045333 | 9 | $6.13{\times}10^{-4}$ |
| Cellular response to hypoxia | GO:0071456 | 7 | $6.23{\times}10^{-4}$ |
| **Cluster E** | | | |
| None | | | |
| **Cluster F** | | | |
| Regulation of protein stability | GO:0031647 | 13 | $7.48{\times}10^{-6}$ |
| Positive regulation of protein catabolic process | GO:0045732 | 10 | $2.07{\times}10^{-5}$ |
| Homeostasis of number of cells | GO:0048872 | 12 | $2.12{\times}10^{-5}$ |
| Regulation of protein catabolic process | GO:0042176 | 13 | $2.51{\times}10^{-5}$ |
| Stress-activated MAPK cascade | GO:0051403 | 10 | $6.01{\times}10^{-5}$ |
| Protein exit from endoplasmic reticulum | GO:0032527 | 5 | $7.06{\times}10^{-5}$ |
| Stress-activated protein kinase signaling cascade | GO:0031098 | 10 | $7.93{\times}10^{-5}$ |
| Regulation of protein exit from endoplasmic reticulum | GO:0070861 | 4 | $9.55{\times}10^{-5}$ |
| Response to ketone | GO:1901654 | 9 | $1.17{\times}10^{-4}$ |
| Regulation of metal ion transport | GO:0010959 | 13 | $1.26{\times}10^{-4}$ |
| **Cluster G** | | | |
| Sterol biosynthetic process | GO:0016126 | 4 | $1.88{\times}10^{-4}$ |
| Eye development | GO:0001654 | 8 | $3.45{\times}10^{-4}$ |
| Visual system development | GO:0150063 | 8 | $3.70{\times}10^{-4}$ |
| Sensory system development | GO:0048880 | 8 | $4.09{\times}10^{-4}$ |
| Cholesterol metabolic process | GO:0008203 | 5 | $4.29{\times}10^{-4}$ |
| Kidney development | GO:0001822 | 7 | $5.03{\times}10^{-4}$ |
| Kidney epithelium development | GO:0072073 | 5 | $5.20{\times}10^{-4}$ |
| Regulation of osteoblast differentiation | GO:0045667 | 5 | $5.37{\times}10^{-4}$ |
| Secondary alcohol metabolic process | GO:1902652 | 5 | $5.88{\times}10^{-4}$ |
| Renal system development | GO:0072001 | 7 | $6.07{\times}10^{-4}$ |
| **Cluster H** | | | |
| Ameboidal-type cell migration | GO:0001667 | 18 | $4.46{\times}10^{-7}$ |
| Epithelial cell migration | GO:0010631 | 15 | $1.13{\times}10^{-6}$ |
| Epithelium migration | GO:0090132 | 15 | $1.25{\times}10^{-6}$ |
| Tissue migration | GO:0090130 | 15 | $1.48{\times}10^{-6}$ |

(*continued on next page*)

**Supplemental Table 1. GO analysis of cluster B-J genes (*continued*)**

| Pathway | GO ID | Gene number | *p*-value |
|---|---|---|---|
| Endothelial cell migration | GO:0043542 | 13 | $1.49\times10^{-6}$ |
| Cellular response to transforming growth factor beta stimulus | GO:0071560 | 12 | $6.23\times10^{-6}$ |
| Response to transforming growth factor beta | GO:0071559 | 12 | $7.77\times10^{-6}$ |
| Regulation of ossification | GO:0030278 | 8 | $1.10\times10^{-5}$ |
| Regulation of epithelial cell migration | GO:0010632 | 12 | $1.31\times10^{-5}$ |
| Retina vasculature development in camera-type eye | GO:0061298 | 4 | $1.65\times10^{-5}$ |
| **Cluster I** | | | |
| Plasma membrane phospholipid scrambling | GO:0017121 | 5 | $1.43\times10^{-6}$ |
| Phospholipid translocation | GO:0045332 | 5 | $5.90\times10^{-5}$ |
| Viral genome replication | GO:0019079 | 7 | $8.09\times10^{-5}$ |
| Positive regulation of epithelial cell proliferation | GO:0050679 | 9 | $8.47\times10^{-5}$ |
| Lipid translocation | GO:0034204 | 5 | $9.20\times10^{-5}$ |
| Regulation of membrane lipid distribution | GO:0097035 | 5 | $1.38\times10^{-4}$ |
| Regulation of epithelial cell proliferation | GO:0050678 | 12 | $1.43\times10^{-4}$ |
| viral process | GO:0016032 | 12 | $1.67\times10^{-4}$ |
| Defense response to virus | GO:0051607 | 10 | $1.68\times10^{-4}$ |
| Semaphorin-plexin signaling pathway involved in axon guidance | GO:1902287 | 3 | $1.72\times10^{-4}$ |
| **Cluster J** | | | |
| Regulation of protein-containing complex disassembly | GO:0043244 | 8 | $7.05\times10^{-5}$ |
| Histone modification | GO:0016570 | 16 | $8.66\times10^{-5}$ |

*Asterisks indicate pathways associated with the regulation of cell cycle and DNA replication in clusters B, C, and D.
